# A nanoscale map of the human cell surface proteome reveals spatial and functional organization

**DOI:** 10.1101/2025.02.12.637979

**Authors:** Brendan M. Floyd, Elizabeth L. Schmidt, Nicholas A. Till, Benjamin S. Waldman, Jonathan L. Yang, Pinyu Liao, Benson M. George, Jan E. Carette, Ryan A. Flynn, Carolyn R. Bertozzi

## Abstract

The cell surface proteome governs processes essential for development, immunity, and disease, yet its large-scale spatial organization remains uncharted. Here, we systematically map the surface proteome of human T-lymphocytes and B-lymphoblasts using proximity labeling of 85 antigens. Coupled with data-independent acquisition mass spectrometry, this approach reveals functional modules, including known complexes like the T-cell receptor alongside clusters not previously implicated in cell membrane functions. Notably, we identify mitochondrial proteins at the surface, suggesting they may bear underappreciated roles at the plasma membrane. A machine learning classifier predicts surface protein associations, revealing IL10RB’s regulation of type I interferon signaling. Spatial modeling distinguishes widely dispersed from localized antigens, indicating active mechanisms of surface organization. In summary, this work provides a large-scale map of the human surfaceome and a resource for exploring its spatial and functional organization.

## Introduction

Proteins located on the plasma membrane and exposed to the extracellular space, collectively known as the surfaceome, are critical for a variety of cellular processes such as immune cell recognition of damaged cells and formation of complex neural circuits^1^. Proteins of the surfaceome provide the primary means of cellular response to extracellular cues and stimulation with different ligands^2,3^. Aberrations in the expression and arrangement of cell surface proteins has been linked to various disease states, including breast^4^, colorectal^5^, and pancreatic cancers^6^. These disease-related changes are often associated with alterations in signaling pathways that depend on protein interactions formed at the surface. The plasma membrane is thought to consist of spatially grouped proteins and lipids that form micro- and nanodomains^7^. These domains are considered hubs for cooperative signaling among protein constituents. To date, studies of functional domains on the cell surface have primarily been conducted in either reconstituted systems that lack native cellular context or in individual units, limiting our understanding of the cell surface as an integrated system. A large-scale map of these functional domains on the cell surface remains uncharted.

Recently developed proximity labeling (PL) proteomics methods have been used to probe cell surface protein-protein interactions^9,10^. These methods deploy chemical probes that are converted to reactive intermediates in the presence of a localized catalyst, typically an enzyme or a photocatalyst. The short half-life of the reactive intermediate limits its diffusion distance, leading to spatially-constrained protein labeling in proximity to the catalyst. Several enzymes have been employed as PL catalysts, including the engineered ascorbate peroxidase APEX2^11^, the relatively promiscuous horseradish peroxidase (HRP)^12^, and biotin ligase variants such as BioID^13^ and TurboID^14^. These methods differ in their labeling radii and reaction times, driving researchers to explore methods with finer control. Photocatalytic methods leverage either Dexter energy transfer (e.g. μMap) or singlet oxygen generators (e.g., the Eosin Y-catalyzed Multimap and the Rhodamine dye-catalyzed LUX-MS)^15–18^. Fluorescence microscopy has also been used to profile the surface proteome, with methods like Förster resonance energy transfer (FRET)^19^ and super-resolution techniques, such as STED^20^, dSTORM^21^, and DNA-PAINT^22^, providing insights into protein clustering and nanodomain organization^23^.

Collectively, these approaches have enhanced our understanding of specific protein clusters. Furthermore, proximity labeling methods have been used to systematically map intracellular protein complexes and functional associations related to disease states^24–26^. By contrast, efforts to define the large-scale interactome and functional associations of the surfaceome have been limited.

Here, we used PL, mass spectrometry and machine learning methods to construct a high-resolution interactome of the cell surface proteomes of human lymphocytes. We targeted 65 surface antigens on T lymphocyte cell lines with PL probes, identified their interactors, and applied unsupervised clustering of the resulting dataset to reveal functional associations on the T cell surface. This analysis identified a class of previously uncharacterized surface antigens, including a subset of proteins conventionally identified as mitochondrial. We further developed a supervised machine learning classifier to predict protein associations across the cell surface proteome. Using this classifier, we identified and characterized a regulatory interaction between the IL-10 receptor beta subunit and the Type 1 Interferon complex. Expanding our analysis, we mapped the cell surface proteome on B lymphoblasts by proximity labeling 20 surface antigens and characterized both conserved protein associations and functional nanodomains specific to these cells. Finally, we used three-dimensional network graphs to model the spatial distribution of surface proteins on T cells and B cells, demonstrating the utility of this dataset for exploring fundamental questions about cell surface biology. This study provides a large-scale spatial analysis of the cell surface proteome in T lymphocytes and B lymphoblasts, in addition to a combination of protocols, open-source code, and a web-accessible data resource (cellsurfacemap.org).

## Results

### Systematic proximity labeling of a T lymphocyte cell surface proteome

To explore the functional organization and spatial distribution of the T-cell surface proteome, we conducted proximity labeling experiments using wild-type Jurkat cells. We hypothesized that the broad labeling radius (200–300 nm^27^) of the horseradish peroxidase (HRP)-catalyzed PL reaction would enable the creation of overlapping nanodomains around target proteins. Thus, targeting a critical mass of surfaceome of components would lead to broad coverage across the entire cell surface proteome and produce a spatial map of nearest neighbors. We labeled select target proteins with primary antibodies, then with a secondary antibody conjugated to HRP (**Figure 1A**). In the presence of hydrogen peroxide, HRP converts biotinyl tyramide into a phenoxyl radical with a short half-life (<1 ms), labeling proteins within a 200-300 nm radius of the target protein. After labeling, cells were lysed, the membrane fraction was isolated, and biotinylated proteins were enriched via streptavidin precipitation. We quantified protein abundances using an optimized data-independent acquisition (DIA) mass spectrometry workflow, incorporating gas-phase fractionation injections to build a comprehensive spectral library^28,29^. Fold changes for all detected proteins were then calculated relative to an isotype control sample.

**Figure 1.**
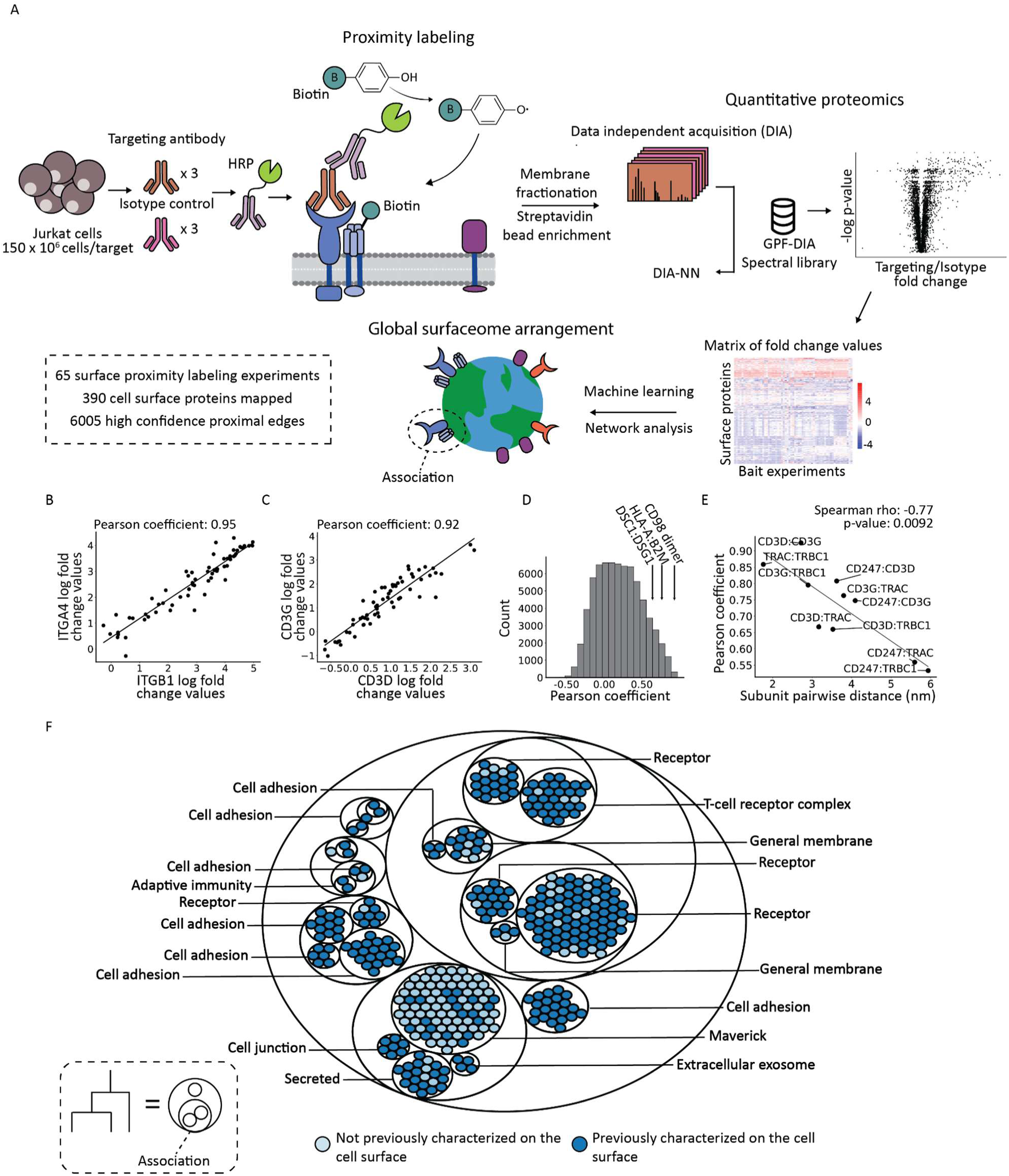
Systematic proximity labeling of the Jurkat cell surface proteome reveals functional clusters on the cell surface. (A) Overview of the approach to spatially map the Jurkat cell surface proteome. DIA-NN: DIA by neural networks; GPF-DIA: Gas phase fractionation data independent acquisition. (B) Scatterplot of log fold change enrichment values and correlation coefficients for the known protein interactors (B) ITGB1/ITGA4 and (C) CD3 complex members CD3D and CD3G. (D) Distribution of pairwise Pearson correlation coefficient across all Jurkat surface protein pairs. Additional known protein interactions are highlighted. (E) Scatterplot of T-cell receptor subunit pairwise distances compared to their estimated Pearson correlation values from Jurkat surface proximity labeling experiments. Spearman correlation p-values were calculated using SciPy’s two-sided test of the null hypothesis that the monotonic association between pairwise subunit distance and Pearson correlation is zero. (F) Circle plot of the clustered Jurkat cell surface proteome labeled with enriched functional annotations for each cluster and colors indicating cell surface characterization of individual surface proteins.

To validate that our workflow accurately enriched neighboring proteins of target proteins, we used an antibody targeting the receptor tyrosine phosphatase CD45, or PTPRC. This experiment enriched 145 surface proteins neighboring CD45. A comparison with a previously published dataset showed an 84% overlap in enriched proteins and fold change values that largely corresponded with each other (**Extended Data 1A, Extended Data 1B**)^14^, demonstrating the reproducibility of this proximity labeling workflow and proteomics approach. Additionally, we employed the high-resolution μMap approach, using an iridium-catalyst conjugated secondary antibody for CD45-targeted PL. We confirmed that proteins enriched with this method also overlapped with those from the HRP-catalyzed biotinylation approach (**Extended Data 1C**). To assess potential antibody-specific biases, we performed proximity labeling of the surface protein ITGB2/CD18 using three independent antibody clones (1B4/CD18, CBR LFA-1/2, and TS1/18) with HRP-based labeling. Across the three clones, 93% of enriched proteins (fold change ≥ 2, adjusted p-value < 0.05) were shared indicating minimal antibody-specific bias (**Extended Data 1D**).

With confidence in our experimental workflow, we systematically mapped the cell surface proteome. To identify potential bait proteins, we used a wheat germ agglutinin horseradish peroxidase (WGA-HRP) conjugate to label the Jurkat cell surface proteome with biotin^30^. Following streptavidin capture, enriched cell membrane proteins were identified using quantitative mass spectrometry. From the resulting list of cell surface proteins, we selected potential bait proteins across a range of protein abundances using the following criteria: (1) previously reported protein function on the cell surface, (2) availability of commercial antibodies against the target protein, and (3) expression of target proteins on other cell types to allow for comparative studies (**Extended Data 1E**). This selection process yielded over 90 surface antigens, for which we screened 94 antibodies for cell surface binding affinity on Jurkat cells. Of these, antibodies against 65 surface antigens showed significant cell surface binding, making them suitable for proximity labeling (**Extended Data 1F, Table S1**). We then applied these antibodies in our optimized workflow, confirming target antigen enrichment to verify antibody specificity. Including replicates, gas-phase fractionation spectral library runs, isotype control samples, and antibody-targeted samples, a total of 339 mass spectrometry experiments were conducted with labeled Jurkat cells. The resulting data identified 390 Jurkat cell surface proteins, in good agreement with a previously reported machine learning predicted surfaceome dataset (i.e., 77% overlap) (**Extended Data 1H**). Overall, our final 65-dimensional dataset contains 25,806 fold change enrichment values that define the spatial organization of the Jurkat cell surface proteome (**Table S2**).

### Prey-centric analysis reveals functional protein modules on the cell surface

Our dataset, encompassing enrichment profiles from each of the proximity labeling experiments, provided a framework for examining the spatial relationships between proteins on the Jurkat cell surface (**Extended Data 1I**). We first sought to verify that we were capturing known protein-protein interactions. We reasoned those proteins located proximally to each other would share similar enrichment values across the 65 proximity labeling experiments. As expected, known protein interactors including the integrin α4β1 heterodimer (**Figure 1B**) and CD3 complex subunits (**Figure 1C**) showed high correlation (Pearson coefficients of 0.95 and 0.92 respectively) across the proximity labeling experiments. Additionally, several other dimeric or multi-subunit complexes exhibited strong correlations across experiments, such as the CD98 heterodimer, the MHC class I complex members HLA-A and B2M, and the cell junction proteins Desmoglein 1 and Desmocollin 1 (**Figure 1D, Extended Data 1I**). These results reinforced the reliability of our surfaceome map in accurately capturing known protein interactions.

Next, we aimed to assess the spatial resolution of our dataset. We benchmarked pairwise Pearson correlation coefficients of the subunits of the CD3-T-cell receptor complex by comparing them to actual distances determined from the center of mass of a 3.7 Å cryo-EM structure (PDB 6JXR)^31^. A significant correlation was observed between the pairwise Pearson coefficients and the true distances between subunits, demonstrating the systematic proximity labeling approach’s ability to discern subunit arrangements down to approximately 4 nm, notably a much higher resolution than the labeling radius of a single experiment (**Figure 1E, Extended Data 1J**).

We then explored whether functional modules could be identified on the cell surface. To achieve this, we used a hierarchical clustering approach previously employed to analyze multimeric protein complexes in plants^32^ and red blood cells^33^. In short, this approach relies on Euclidean distance-based clustering of proteins using their enrichment profiles in the 65-dimensional dataset. Rather than applying a single cutoff to define hierarchies, multiple cutoffs are used to define discrete modules of spatially co-localized proteins^32^. For each cluster, we identified enriched functional annotations using the DAVID web server^34,35^ and determined whether the surface antigen had been previously characterized on the cell surface (**Figure 1F, Extended Data 1K, Table S2**). This analysis uncovered several functional modules, with a notable presence of cell adhesion-related clusters. Multi-subunit complexes, including the T-cell receptor alpha and beta subunits along with CD3 complex members, clustered with other adaptive immunity regulators, such as the leukocyte associated inhibitory receptor LAIR1 and the MHC class 1 subunits HLA-A and HLA-B. Interestingly, cell junction associated proteins normally involved in intercellular adhesion such as Desmoglein 1 and Desmocollin 1 clustered closely with proteins associated with secreted vesicles and extracellular exosomes, possibly pointing to a location of material exchange between cells. In all, this analysis grants a high-level view of the functional compartmentalization on the cell surface.

### Conservation of cell surface protein organization across cell types

To explore differences and similarities in surface protein arrangement across cell types, we adapted our proximity labeling and surface mapping approach for Daudi cells, a B-lymphoblast line. Daudi cells offer a distinct perspective as they express B-cell-specific proteins like CD19, CD20, and CD22, while also sharing some surface markers with Jurkat cells, allowing for comparative analysis. We proximity labeled 20 surface antigens on Daudi cells, including nine B-cell-specific antigens and 11 antigens shared with Jurkat T-lymphocytes (**Figure 2A, Extended Data 2A, Table S3**). Using criteria similar to those applied to our Jurkat dataset but using a lower experimental enrichment threshold for inclusion due to the smaller amount of Daudi proximity labeling experiments, we identified and mapped 189 proteins on the Daudi cell surface (**Extended Data 2B, Extended Data 2C**). We then applied a prey-centric analysis to the Daudi cell surface proteome proximity labeling dataset and for validation we examined the correlation of known protein interactors across runs. Among the most correlated pairs were the two subunits of the Human Leukocyte Antigen DR isotype (HLA-DR) complex (**Figure 2B**). Additionally, the Igα (CD79A) and Igβ (CD79B) subunits of the B-cell receptor complex demonstrated high pairwise correlations (**Figure 2C**, **Extended Data 2D**).

**Figure 2.**
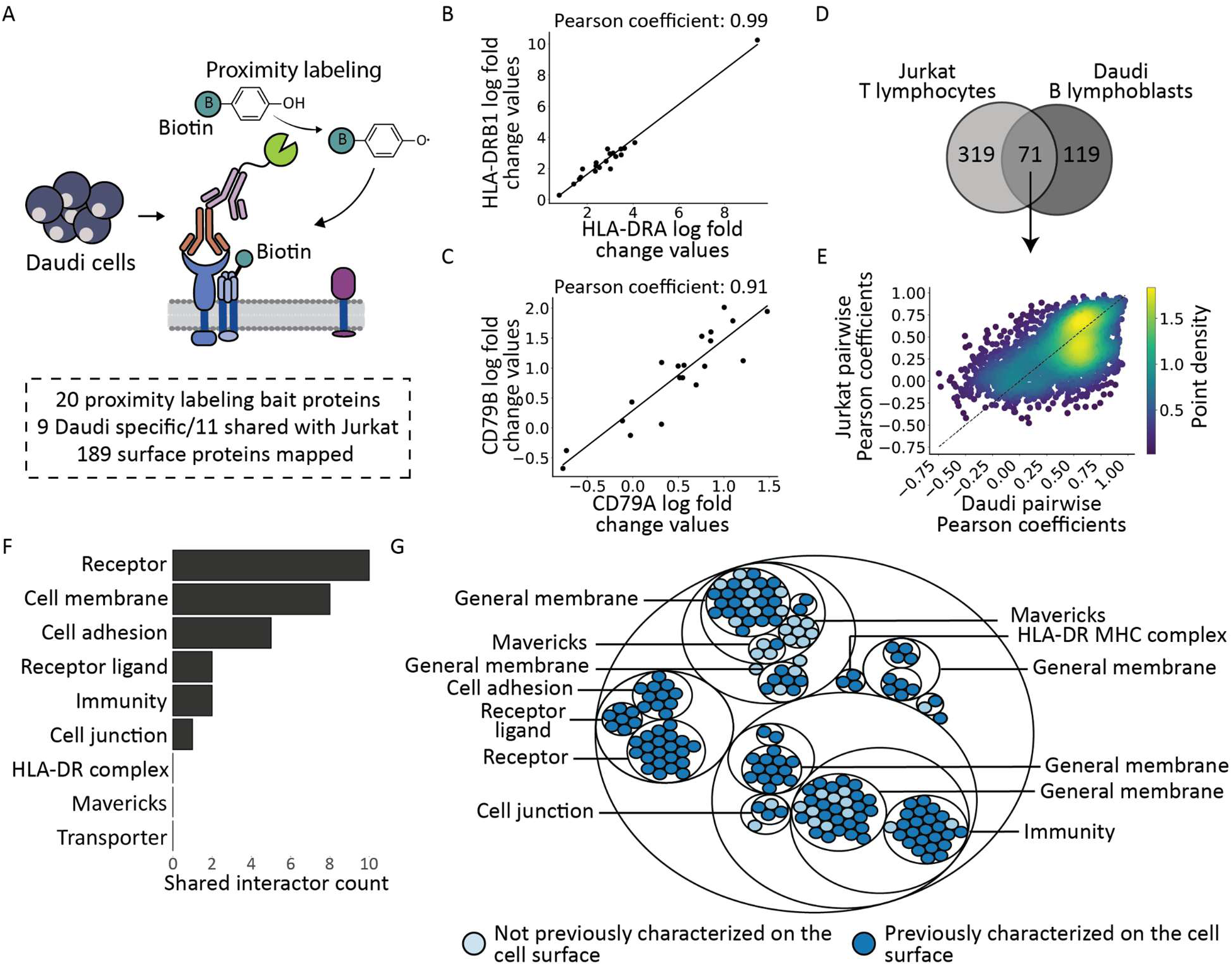
Cell surface protein interactions are conserved across different cell types. (A) Numerical summary of cell surface proximity labeling done on Daudi B-lymphoblast cells. (B) Scatterplot of log fold change enrichment values and correlation coefficients for the MHC class 2 subunits HLA-DRA and HLA-DRB1 (C) and B-cell receptor SRK subunits CD79A and CD79B. (D) Venn diagram showing overlap of cell surface proteins identified on Jurkat and Daudi cells. (E) Scatterplot of pairwise Pearson correlation coefficients in Daudi and Jurkat cell lines. The dashed line represents a line of perfect correspondence where *y = x*. (F) Functional annotation counts of conserved protein-protein interactions in Daudi and Jurkat cell lines. Conserved protein-protein interactions are defined as protein pairs with Pearson coefficients greater than 0.7 and difference between the two cell lines of less than 0.1. (G) Circle plot of the clustered Daudi cell surface proteome labeled with enriched functional annotations for each cluster and colors indicating cell surface characterization of individual surface proteins.

Next, we analyzed the overall similarities in pairwise protein interactions between Jurkat and Daudi cells. From the cell surface proteomes of both cell types, we identified 71 shared proteins (**Figure 2D**). These proteins included functional units essential for various immune cell types, such as cell adhesion and immune-related proteins. Using this subset of shared proteins, we calculated pairwise Pearson correlation values for all protein pairs and compared them between the two cell lines (**Figure 2E**). This comparison revealed strong correspondence between the 2,278 pairwise associations, with a significant Spearman rho value of 0.516. Focusing on interactions with nearly identical Pearson values (within 0.05) between the two cell types, we observed a striking prevalence of receptor interactions (**Figure 2F**). Finally, we clustered the Daudi cell surface proteome, highlighting functionally enriched annotations for each cluster (**Figure 2G, Extended Data 2E**). This clustering captured a cell junction protein cluster similar to that observed on Jurkat cells in addition to a cluster of immunity-related proteins, including the B-cell receptor accessory proteins CD72 and CD19. Notably, the MHC class II HLA-DR complex subunits formed their own distinct cluster.

### Discovery of clusters of uncharacterized cell surface mitochondrial proteins

Among the identified and clustered surface proteins on Jurkat cells, 72% had been previously characterized on the cell surface. However, we observed a distinct cluster of proteins typically assumed to be restricted to mitochondria. These mitochondrial proteins constitute approximately 10% of our observed surface proteins on Jurkat cells (**Figure 3A**). We also observed chaperone proteins such as the endoplasmic reticulum-localized HSP90B1^36^ and RNA binding proteins such as the nuclear RNA binding-protein FUS^22^. We refer to these non-canonical cell surface proteins as “maverick proteins”.

**Figure 3.**
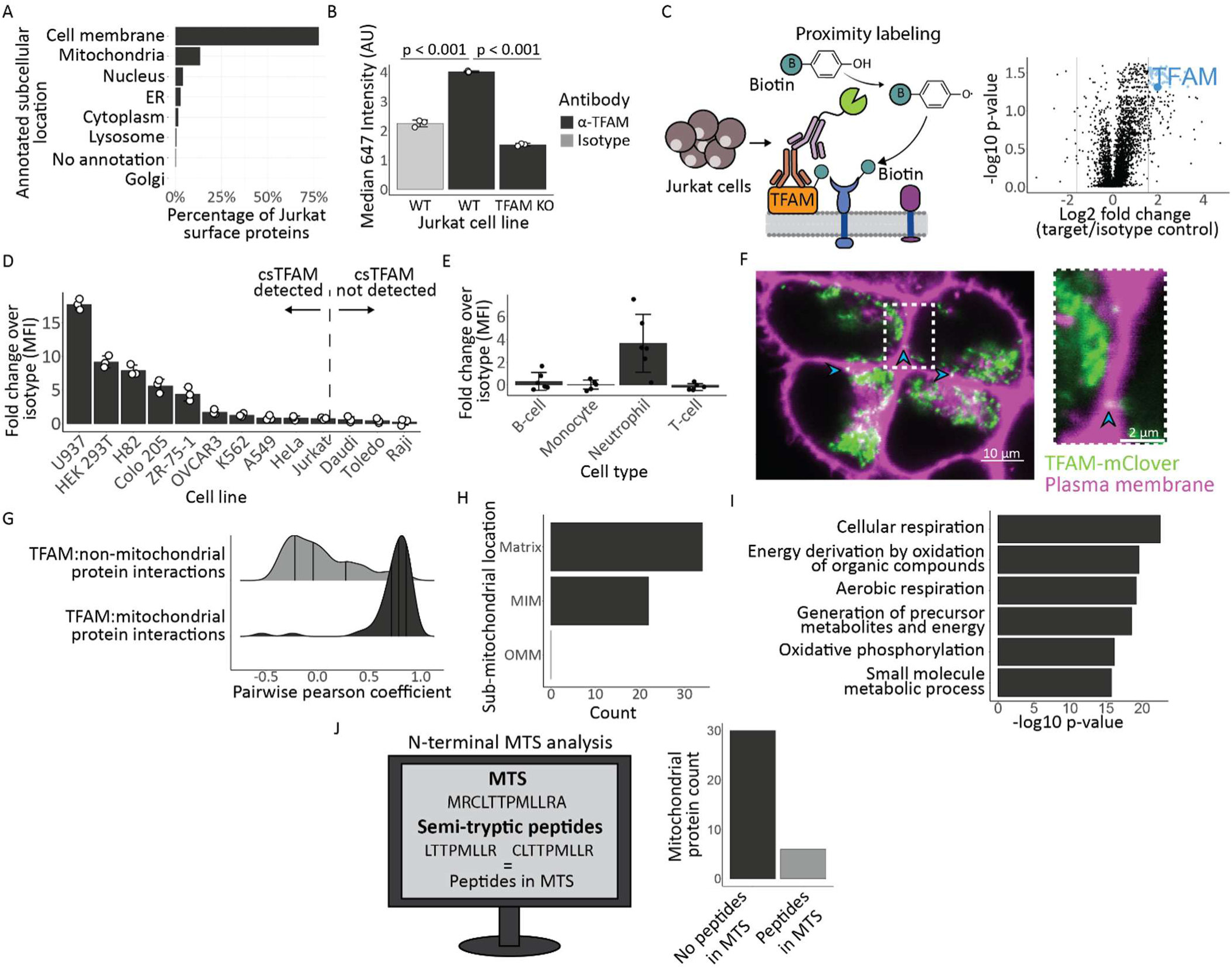
Detection and validation of maverick mitochondrial proteins on the cell surface of Jurkat cells. (A) Breakdown of annotated subcellular localizations for the Jurkat cell surface proteome. (B) Flow cytometry cell surface staining with an isotype control (grey) and anti-TFAM (black) antibody on Jurkat WT and TFAM KO cell lines. P-values were estimated using a two-sided Student’s T-test with p-values of 1.3878e-5 and 1.288e-7 and t-statistics of 25.58 and -82.6013 for n=3 biological replicates. Error bars represent the standard deviation from the mean. (C) Cartoon of the cell surface proximity labeling experiment using TFAM as a bait. Volcano plot showing the results of cell surface proximity labeling using an anti-TFAM antibody. TFAM is highlighted in dark blue and other enriched proteins are shown in light blue. P-values were estimated using a two-sided t-test with false discovery rate (FDR) correction performed using permutation-based FDR with 250 randomizations for n = 3 biological replicates. (D) Anti-TFAM flow cytometry cell surface staining results across 13 different cell lines. Fold change was estimated by comparing to isotype control binding levels on each cell line and error bars correspond to standard deviation from the mean for n = 3 biological replicates. (E) Anti-TFAM cell surface staining results for different cell types from 6 different donors. Fold change was estimated by comparing to isotype control binding levels and error bars correspond to standard deviation from the mean. (F) Confocal microscopy image of HEK-293T cells expressing a TFAM-mClover fusion and stained with CellMask Orange plasma membrane stain. The dashed box indicates the region shown at higher magnification. Arrowheads denote regions of TFAM-mClover localization at the plasma membrane. mClover fluorescence was detected in the 488 nm channel and the plasma membrane stain in the 561 nm channel. Confocal microscopy of HEK-293T cells expressing a TFAM-mClover fusion protein and stained with CellMask Orange to label the plasma membrane. The dashed box indicates the region shown at higher magnification. Arrowheads highlight regions of TFAM-mClover localization at the plasma membrane. mClover fluorescence was detected in the 488 nm channel, and the plasma membrane stain in the 561 nm channel. This image is representative of an experiment that was repeated 3 times. (G) Distribution of TFAM pairwise Pearson values with other detected surface mitochondrial proteins (grey) and all non-mitochondrial surface proteins (black). (H) Sub-mitochondrial counts for mitochondrial proteins detected on the cell surface. (I) Enriched GO biological process terms for cell surface mitochondrial proteins. A one-tailed cumulative hypergeometric test was used to estimate p-values with a set counts and sizes correction method for false discovery rate control. (J) Description of the analysis done to identify native N-termini peptides from the mitochondrial transit sequence for mitochondrial proteins detected on the cell surface and bar plot showing the count of maverick mitochondrial proteins with peptides detected in the TPPRED3 predicted mitochondrial transit sequence (MTS).

We confirmed that maverick proteins are indeed present on live Jurkat cell surfaces using flow cytometry. We confirmed cell surface expression of the mitochondrial protein transcription factor A (TFAM), which exhibited substantial peptide coverage in our proximity labeling experiments (**Extended Data 3A**), as well as the nuclear RNA-binding protein FUS which was observed both on Jurkat and Daudi cells^22^ (**Extended Data 3B, Extended Data 3C**). To confirm antibody specificity for TFAM, we generated a Jurkat TFAM knockout line using CRISPR-Cas9 (**Extended Data 3D**) and again probed for cell surface TFAM using flow cytometry. As expected, the knockout line showed diminished TFAM antibody binding (**Figure 3B, Extended Data 3E, Extended Data 3F**). Further, we profiled the interactome of cell surface TFAM (csTFAM) via the HRP-mediated PL method (**Figure 3C**). We observed enrichment of 58 proteins (relative to an isotype control), which included TFAM itself as well as other maverick proteins such as FUS and another RNA-binding protein, CIRBP (**Table S2**). We then extended this analysis to 13 additional cell lines as well as human primary peripheral blood mononuclear cells (PBMCs). Cell surface TFAM expression was observed in several cell lines and was particularly high in the U-937 monocytic cancer line (**Figure 3D**). We also detected cell surface TFAM on neutrophils from human PBMCs (**Figure 3E**). To probe the possibility of the detection of csTFAM resulting from the passive adsorption of the protein from dead cells sticking to the membrane we generated a TFAM-mClover3 fusion and expressed it in HEK-293T cell lines. Confocal imaging revealed distinct foci of TFAM localization at the plasma membrane, particularly at points of cell-cell contact (**Figure 3F, Extended Data 3G**).

Having identified TFAM as a *bona fide* surface antigen, we sought to understand its spatial relationship to other mitochondrial maverick proteins. We observed high pairwise correlation values among TFAM and these other proteins, particularly with ATP synthase subunits, aligning with prior reports of ATP synthase translocation to the plasma membrane in cancer cell lines (**Figure 3G**)^37,38^. Notably, all detected mitochondrial proteins on the Jurkat cell surface originated from the mitochondrial inner membrane (MIM) or matrix, with none from the outer membrane (**Figure 3H**). GO enrichment analysis of this cluster highlighted significant roles in cellular respiration, suggesting a potential adaptation for supplemental energy acquisition as previously observed with translocated cell surface ATP synthase^37^ (**Figure 3I**).

The pathway by which mitochondrial proteins traffic to the cell surface is unclear. However, our analysis suggests that they first transit through mitochondria. We predicted mitochondrial targeting sequences (MTS)^39^ for this set of surface-localized mitochondrial proteins and reprocessed a subset of our proximity labeling data, allowing semi-tryptic N-termini to capture proteolytically processed peptides. For proteins with high-confidence MTS predictions, 83% showed no detectable peptides derived from the MTS region, implying they had undergone mitochondrial processing before reaching the plasma membrane (**Figure 3J**). This class of maverick cell surface proteins represents a previously uncharacterized category of surface antigens, with potential applications in cancer targeting as previously demonstrated with ATP synthase^40–42^.

### A machine learning classifier for prioritizing cell surface protein associations from proximity labeling data

In order to discover previously uncharacterized protein functions based on cell surface protein associations, we developed a machine learning approach to prioritize protein pairs with interaction-like proximity labeling signatures. Specifically, we trained a random forest classifier to assign association scores to protein pairs across the Jurkat cell surface proteome based on our proximity labeling data. A random forest classifier was chosen due to its ability to handle high-dimensional data, robustness against overfitting, and interpretability of its feature importance rankings, which are critical for biological datasets with complex data structures^43^. To evaluate whether random forest was appropriate for this prioritization task, we compared its performance in prioritizing known protein-protein interactions (PPIs) when trained on our proximity labeling dataset against eight additional supervised machine learning classifiers, including gradient boosting, XGBoost, ExtraTrees, and linear support vector machine models. The random forest classifier outperformed the other tested models when evaluated by stratified 5-fold cross-validation, supporting its use for prioritizing protein associations from proximity labeling data (**Extended Data 4A)**.

For classifier training, we used binary protein-protein interaction data from multiple sources to compile a set of gold-standard PPIs. From this set, 1,532 PPIs comprised pairs of proteins that were both identified on the surface of Jurkat cells. To prepare the proximity labeling data for classification, we calculated fold change differences and similarity metrics (e.g., Pearson correlation, Euclidean distance) for all possible protein pairs, creating a comprehensive feature matrix for the 60,031 possible protein pairs (**Figure 4A**). We then trained our random forest classifier on this feature matrix using the gold-standard cell surface PPIs as positive labels. With further optimization of the classifier’s hyperparameters, we generated a classifier that robustly prioritized gold-standard protein associations in stratified 5-fold cross-validation, achieving an average ROC AUC of 0.92 (**Figure 4B**) and a precision-recall AUC of 0.62 (**Figure 4C, Extended Data 4B**). We also constructed a more conservative gold-standard PPI set using UniProt-derived PPIs and BioGRID multi-validation physical interaction pairs. Although this more stringent gold-standard definition reduced the number of positive Jurkat cell surface PPI labels from 1,532 to 773, classifier performance and output were largely preserved (**Extended Data 4C**). Association scores from the original and stringent classifiers were significantly correlated (**Extended Data 4D**), and the top 10% highest-scoring protein pairs showed substantial overlap (**Extended Data 4E**), indicating that the classifier output was robust to the gold-standard reconstruction.

**Figure 4.**
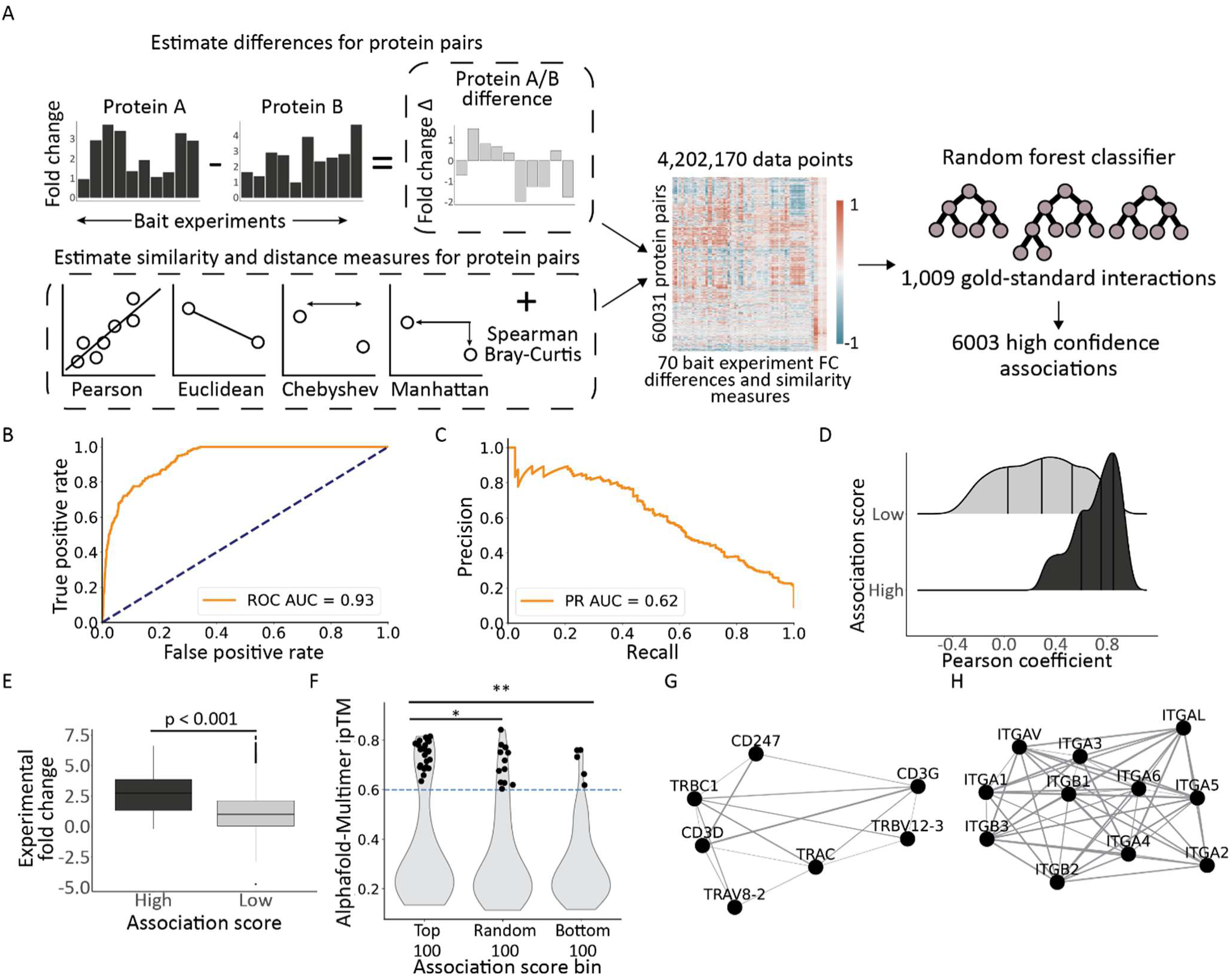
Design of a machine learning classifier for prioritizing cell surface protein associations from proximity labeling data. (A) Workflow for training a random forest classifier to identify cell surface associations. (B) Receiver operator characteristic (ROC) curve for 303 withheld gold standard protein pairs. Area under the ROC curve (ROC AUC) is 0.93. (C) Precision-recall curve for 303 withheld protein pairs. Area under the precision recall curve (PR AUC) is 0.62. (D) Distribution of Pearson correlation coefficients for protein pairs with high random forest classifier association predictions and low random forest classifier predictions. High association scores are defined as any protein pair with a score higher than 0.6. (E) Comparison of experimentally measured proximity labeling fold change (FC) enrichments for 49 bait experiments to the classifier predicted association scores of bait-prey interactions. Two-sided Welch’s T-test statistic 16.69, and p-value 8.045e-49. The box is centered on the median value and the whiskers extend to the smallest and largest data points within 1.5 times the interquartile range from the first and third quartiles, respectively. (F) AlphaFold2-Multimer ipTM values for heterodimer models from the top 100, random 100, and bottom 100 protein pairs based on random forest association scores, with high-ipTM models defined as ipTM ≥ 0.6 and pairwise comparisons performed using a one-sided Fisher’s exact test for top 100 vs random 100 (odds ratio = 2.00, p = 0.0484) and top 100 vs bottom 100 (odds ratio = 5.68, p = 0.000184). (G) Kawada-Kawai network of classifier predictions for T-cell receptor subunits. Edge weight corresponds to association scores. (H) Kawada-Kawai network of classifier predictions for integrin alpha and beta proteins detected on the Jurkat cell surface. Edge weight corresponds to association scores.

We then predicted cell surface associations across the proteome by putting the full set of all-by-all protein pairs through the classifier, which resulted in 6,003 high confidence cell surface protein association predictions (**Extended Data 4F, Table S2**). Because several similarity and distance metrics capture related mathematical properties, we next tested whether these features introduced redundancy that negatively impacted classifier performance. We systematically ablated similarity features prior to classifier training and measured the effect on precision-recall performance (**Extended Data 4G**). The largest drop in average precision was observed when all similarity measures were removed, whereas removal of individual similarity features had more modest effects.

To validate the classifier predictions, we compared the predicted association scores to the estimated Pearson correlation coefficients and observed that higher predicted association scores corresponded to stronger Pearson correlations (**Figure 4D**). As well, comparing our random forest predicted association scores for bait-prey pairwise interactions to the experimentally measured fold change enrichment for our 65 different baits, higher association scores had significantly higher log fold change values compared to interactions with a low association score (**Figure 4E**). For further validation, we asked whether protein pairs with prior interaction annotations received higher classifier association scores. We divided all scored protein pairs into two groups: pairs present in our curated gold-standard PPI set and pairs without a known interaction annotation in that set. Known interaction pairs were strongly enriched for higher classifier scores compared with unannotated pairs (**Extended Data 4H**). Finally, we asked whether high-scoring classifier predictions were enriched for protein pairs with structural support for direct extracellular interactions. To test this, we used AlphaFold2-Multimer^59^ to predict extracellular domain complex structures for three groups of protein pairs: the top 100 highest-scoring random forest predictions, the bottom 100 lowest-scoring predictions, and 100 randomly selected protein pairs. Because proximity labeling can capture direct binding, co-complex membership, and spatial co-localization, we did not expect all high-scoring protein pairs to produce high-confidence structural models. Instead, we tested whether high-scoring random forest predictions were enriched for higher-confidence interface predictions, defined by ipTM ≥ 0.6. Protein pairs in the top-scoring set were significantly enriched for ipTM ≥ 0.6 models compared with both randomly selected pairs and bottom-scoring pairs (**Figure 4F, Table S6**).

Confident that our machine learning classifier was providing high accuracy association predictions, we then looked at known and previously uncharacterized associations predicted for the Jurkat proteome. Known cell surface interactions included those of the TCR subunits (**Figure 4G**). Additionally, we observed high connectivity of the integrin alpha and beta subunits (**Figure 4H**). Because curated protein interaction databases were used to define positive training labels, enrichment of known interaction pairs among high scoring predictions should be interpreted as evidence that the classifier recovers features consistent with existing interaction knowledge, rather than as fully independent validation of uncharacterized association discovery. The ability of this approach to prioritize previously uncharacterized cell surface protein associations therefore requires orthogonal experimental or structural follow up.

### Prediction and characterization of IL10RB as a negative regulator of Type I Interferon signaling

Among the 6,003 high confidence associations predicted by our machine learning classifier, the interaction between the beta subunit of the IL-10 receptor complex, IL10RB, and a subunit of the Type I interferon system, IFNAR2, stood out. IL10RB was predicted to associate with IFNAR2 with an association score of 0.82, placing it in the top 2% of all scored associations (**Figure 5A**). This prediction was particularly intriguing, as IL10RB is a known signaling component for various class II cytokines, including IL-22, IL-26, IL-28, and Type III interferons^44,45^. However, it has not previously been associated with the Type I interferon system. While transcripts for IL10RB co-receptors, such as IL22RA1 and IFNLR1, have been previously detected in Jurkat cells^46^, no protein-level evidence of their translation was detected by mass spectrometry in both whole cell and surface protein enriched samples (**Extended Data 5A**). The lack of canonical interaction partners for IL10RB on the Jurkat cell surface, coupled with its broad involvement in signaling pathways, prompted us to investigate further the classifier prediction of its association with IFNAR2. Interestingly, IL10RB also shares structural similarities with IFNAR1, another core component of the Type I interferon receptor complex including, importantly, an intracellular binding domain for the Jak kinase family member TYK2 (**Extended Data 5B, Extended Data 5C**)^47,48^. To determine if this predicted association is a stable interaction we used AlphaFold2-Multimer and found only a low confidence complex prediction (**Extended Data 5D**).

**Figure 5.**
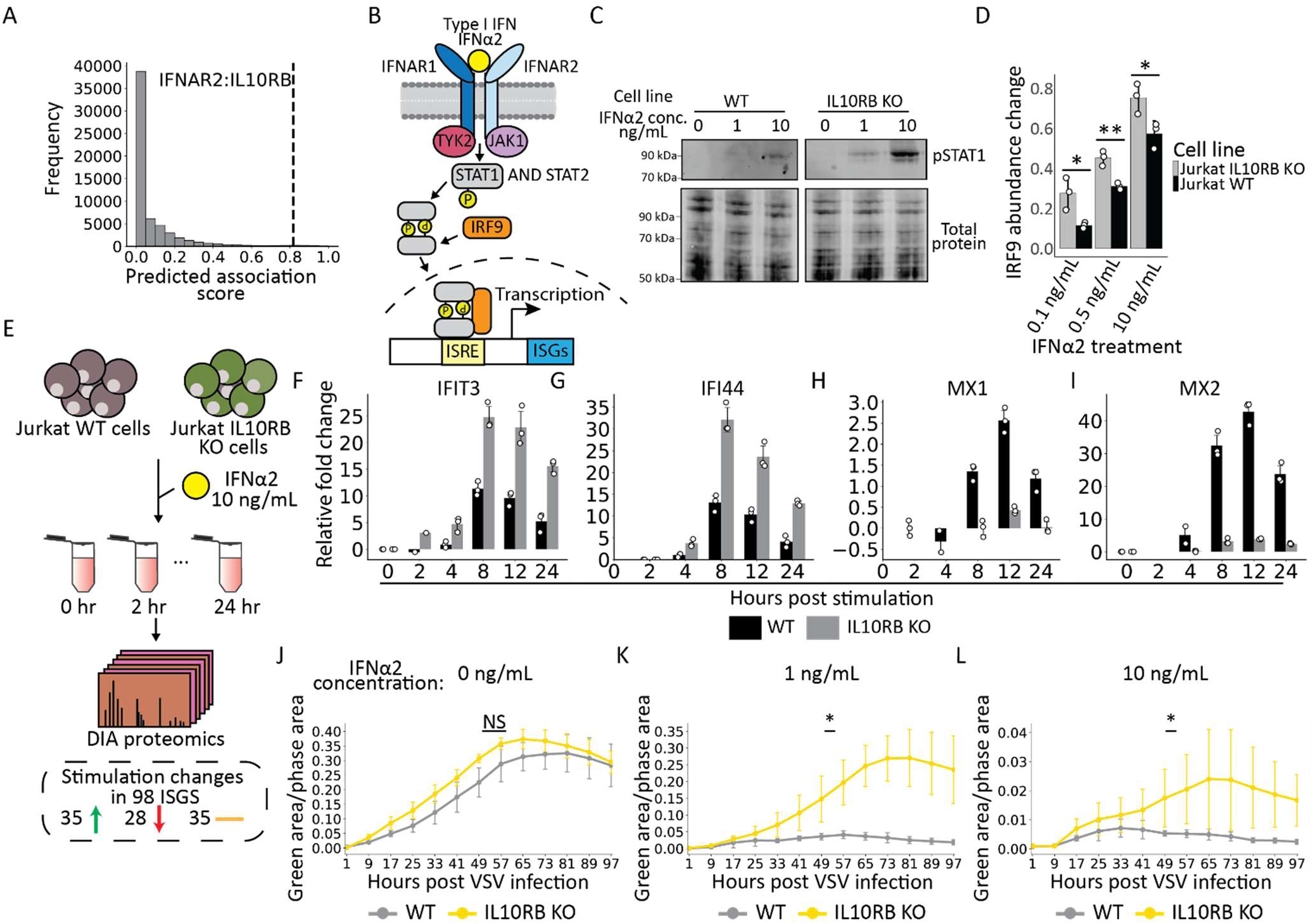
Prediction and characterization of IL10RB as a Type 1 Interferon regulator. (A) Distribution of classifier predicted association scores for all protein pairs with the prediction for IFNAR2 and IL10RB highlighted. (B) Cartoon of the Type 1 Interferon signaling transduction pathway. (C) Western blot of phosphorylated STAT1 (pSTAT1) abundance in IFNα2 stimulated Jurkat WT and IL10RB knockout lines at different interferon concentrations. (D) Intracellular flow cytometry measurement of cytosolic IRF9 abundance change in IFNα2 stimulated Jurkat wild type (black) and IL10RB knockout lines (grey) at various IFNα2 concentrations. Two-sided Student’s t-test p-values for the different concentrations are 0.027, 0.0036, and 0.035 with t-statistics -3.40, -6.12, and -3.13 for 0.1, 0.5, and 10 ng/mL respectively for n=3 biological replicates. Error bars correspond to standard deviation from the mean (E) Cartoon workflow for monitoring interferon stimulated gene protein stimulation using quantitative proteomics when Jurkat WT and IL10RB KO cells were treated with 10 ng/mL of IFNα2 for 0, 2, 4, 8, 12, and 24 h. (F-I) Abundance change plots of selected ISGs (F) IFIT3, (G) IFI44, (H) MX1, and (I) MX2 in the Jurkat WT (black) and IL10RB KO (grey) cell lines. Relative fold change was estimated in comparison to the 0 h time point or the earliest detected time point, and error bars correspond to standard deviation from the mean. N = 3 biological replicates. (J-L) Viral replication curves for Jurkat WT and IL10RB KO cells pre-stimulated with (J) 0, (K) 1, and (L) 10 ng/mL IFNα2. A two-sided Welch’s t-test at the 81-minute timepoint yielded p-values of 0.44, 0.0008, and 0.024, with corresponding t-statistics of −0.81, −6.98, and −3.19 for 0, 1, and 10 ng/mL IFNα2, respectively. Error bars represent standard deviation from the mean for n = 6 biological replicates.

To explore the functional relevance of the IL10RB-IFNAR2 association, we generated an IL10RB knockout (KO) Jurkat cell line using CRISPR-Cas9. As IL10RB and IFNAR2 are located in cis on chromosome 21, we verified that IFNAR2 expression remained intact in the knockout line (**Extended Data 5E**). Type I interferon signaling is mediated through the JAK1/STAT pathway and involves STAT1 and STAT2 phosphorylation, leading to heterodimerization and complex formation with IRF9, which then promotes transcription and activation of interferon-stimulated genes (ISGs) (**Figure 5B**). In the IL10RB KO line, we observed significantly increased levels of phosphorylated STAT1 compared to the wild-type (WT) line when stimulated with the Type I interferon IFNɑ2 (**Figure 5C**). This was corroborated by intracellular flow cytometry, which showed elevated IRF9 levels following 12-hour stimulation with IFNɑ2 (**Figure 5D**). To confirm the specificity of this effect, we generated a T-cell receptor alpha subunit knockout line (TCR-ɑ KO), which was not predicted to associate with the Type I interferon system (**Extended Data 5F, Extended Data 5G**). When stimulated with IFNɑ2 the TCR-ɑ KO line exhibited no changes in interferon signaling relative to WT cells when monitoring IRF9 levels using intracellular flow cytometry, underscoring the specificity of the IL10RB-IFNAR2 interaction (**Extended Data 5H**).

We then evaluated the dynamics of ISG stimulation in the IL10RB KO line using time-course proteomics. Upon stimulation with 10 ng/mL IFNɑ2, we monitored protein abundance changes in WT and IL10RB KO cells using DIA proteomics (**Figure 5E, Table S4**). Of 98 well-sampled ISGs, 35 exhibited increased stimulation rates in the IL10RB KO line including IFIT3, IFI44, XAF1, and DDX60 (**Figure 5F–I**). Notably, however, 28 ISGs including the interferon-induced GTPases MX1 and MX2 showed significantly reduced stimulation in IL10RB KO cells. (**Extended Data 5I**)^49^. These findings indicate that loss of IL10RB does not uniformly amplify the interferon response but instead alters the composition of the ISG program.

To assess the functional consequences of this altered ISG landscape, we evaluated viral replication in WT and IL10RB KO Jurkat cells using a vesicular stomatitis virus (VSV-GFP) replication assay. Cells were pre-stimulated with IFNɑ2 for 24 hours to establish an interferon-induced antiviral state prior to infection and then infected with VSV-GFP. Despite exhibiting enhanced interferon signaling and increased induction of many ISGs, IL10RB KO cells displayed significantly increased viral replication compared to WT cells across interferon doses (**Figure 5I–K**). Given the established antiviral role of MX1 against VSV, its reduced induction in the IL10RB KO line is likely a major contributor to this phenotype^57,58^. These findings suggest that IL10RB loss results in a qualitatively altered interferon response in which heightened signaling amplitude and induction of select ISGs does not translate into effective viral restriction. Instead, IL10RB appears to be required for proper coordination of antiviral effector expression downstream of Type I interferon signaling, and its absence leads to a dysregulated ISG program that is permissive to viral replication.

### Estimation of cell surface distribution of the cell surface proteome

We next investigated whether our proximity labeling dataset could be used to construct a virtual image of the surface distribution of individual proteins. Such visualization provides insights into the spatial organization of the cell membrane and the mechanisms governing surface protein distributions. Inspired by recent applications of DNA nanotechnology for mapping the spatial arrangement of a limited number of surface proteins^51^, we constructed a Kamada-Kawai spring network based on enrichment connections among 48 high-quality bait proteins on Jurkat cells. The network coordinates were transformed into a three-dimensional (3D) spherical representation, enabling us to project the normalized fold change of a protein of interest across the bait network onto this 3D model. This approach effectively models the distribution of proteins across the cell surface proteome using a metric we term the graph connectivity score. Applying this method to the entire Jurkat cell surface proteome, we estimated the surface coverage of each protein by calculating its relative abundance across the modeled cell surface (**Figure 6A**).

**Figure 6.**
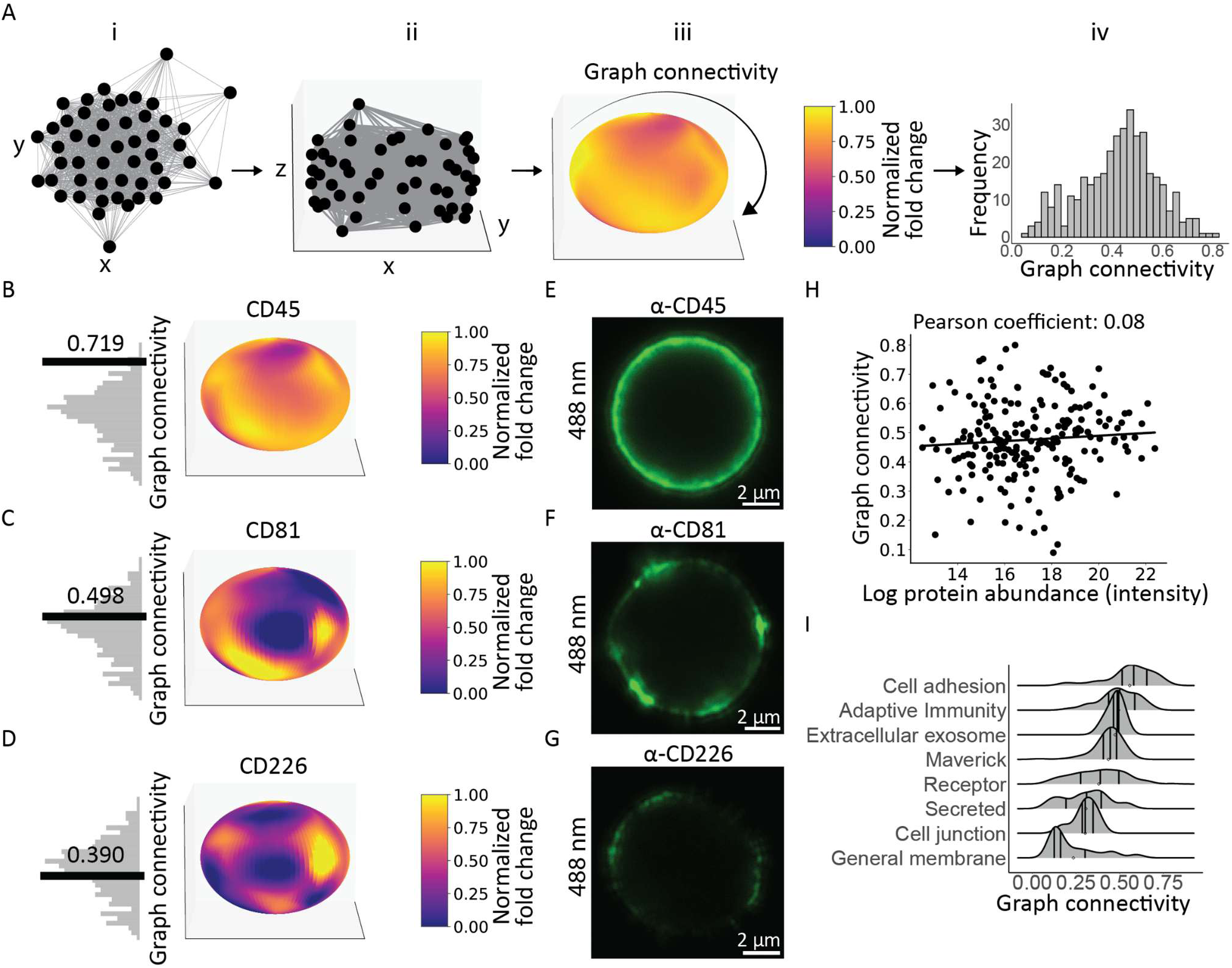
Three-dimensional network modeling to predict the surface distribution of the cell surface proteome. (A) Design walkthrough of 3D network heatmaps where (i) a 2D Kamada-Kawaii network of high-quality proximity labeling bait proteins are (ii) transformed to lie on a 3D sphere where (iii) the normalized fold change of a given surface protein of interest is projected for each bait experiment (iv) and graph connectivity can be estimated for the entire cell surface proteome. (B) 3D network heatmap models of CD45, (C) CD81, and (D) CD226. (E-G) Representative confocal microscopy images of immunostained cell surface antigens CD45 (E), CD81 (F), and CD226 (G). Antigens were visualized in the GFP channel using the dye Alexa Fluor 488. Images are representative of an experiment that was independently repeated 3 times. (H) Scatterplot comparing the protein abundance of Jurkat cell surface proteins measured using WGA-HRP for surface protein enrichment compared to their estimated graph connectivity. (I) Distribution of estimated graph connectivity for cell surface proteins based on their functional annotation.

To quantify protein distribution, we developed the connectivity score, a metric that integrates a protein’s enrichment across the bait network (**Table S2**). Coverage scores varied widely across the proteome, with proteins such as the pan-lymphocyte marker CD45 (coverage score = 0.719) exhibiting widespread distribution, while others, like the tetraspanin web-associated protein CD53 (coverage score = 0.227), displayed more restricted distributions. To determine whether our graph connectivity score was accurately predicting protein cell surface distribution we employed immunofluorescence microscopy to image the distribution of antigens on the cell surface. We first modeled the surface proteins CD45 (**Figure 6B**), CD81 (**Figure 6C**), and CD226 (**Figure 6D**), which represent a range of connectivity scores (**Movie S1)**. Using confocal microscopy, we then imaged the distribution of these surface antigens and after staining cells with antibodies for these antigens, we measured median fluorescence intensity along the cell periphery to estimate their surface distributions. The microscopy data for CD45 (**Figure 6E**, **Extended Data 6A**), CD81 (**Figure 6F**, **Extended Data 6B**), and CD226 (**Figure 6G**, **Extended Data 6C**) closely aligned with the predicted surface distributions, confirming the reliability of our coverage score metric (**Extended Data 6D**). It is worth noting that the graph connectivity score was independent of the protein abundance measured in surface proteomics experiments (**Figure 6H**).

As a practical application of this method, we explored the relationship between surface protein distribution and function. Cell adhesion-related proteins were broadly distributed across the surface, consistent with their role in mediating multiple attachment points. In contrast, receptors demonstrated a more restricted distribution (**Figure 6I**). Interestingly, cell junction proteins showed reduced dispersion relative to adhesion proteins, suggesting distinct organizational features. This pattern was conserved in Daudi cells, highlighting a shared aspect of surfaceome organization (**Extended Data 6E**).

## Discussion

In this work, we systematically mapped the surfaceome of two distinct cell types. By employing a low-resolution proximity labeling approach across 65 surface antigens on T-lymphocytes and 20 antigens on B-lymphoblasts, we assembled a high-resolution interactome for the surfaceome (**Table S5**). While the cell membrane is a dynamic environment, and our dataset represents an averaged snapshot that does not fully capture temporal or state-dependent variability, it nonetheless provides valuable insights into surfaceome organization. From these interactomes, we identified functional modules on the cell surface that encompassed both known protein assemblies, such as the T-cell receptor (TCR) and HLA class I and II complexes, as well as previously uncharacterized clusters.

One particularly intriguing finding was the identification of clusters of maverick surface antigens, including proteins typically localized to intracellular organelles. In Jurkat cells, these were primarily mitochondrial proteins, such as the transcription factor TFAM. Reports of mitochondrial proteins, including the ATP synthase complex, maintaining functionality on the plasma membrane suggest a broader phenomenon of mitochondrial-to-plasma membrane translocation^37,38,40^. Recent studies describing vesicles derived from the mitochondrial inner membrane, which contain TFAM and other mitochondrial proteins, offer a plausible pathway for their delivery to the cell surface even under homeostatic conditions^52^. Additionally, questions remain regarding how mitochondrial matrix proteins lacking transmembrane domains are tethered to the cell surface, whether via direct interactions with canonical transmembrane proteins^53^, lipid-anchored proteins, or other attachment mechanisms^54^. Beyond mitochondrial proteins, the recent characterization of RNA-binding proteins on the cell surface, along with the clustering observed among them, suggests a potential conserved role for these proteins in cellular interactions and signaling^22^. These findings highlight an exciting avenue for future research into both mitochondrial and RNA-binding protein dynamics and their functions at the cell surface.

We developed a high-accuracy machine learning classifier to prioritize protein-protein associations on the cell surface from our proximity labeling data. While this classifier is adaptable to other proximity labeling datasets, its application to different cell lines or tissues would require the assembly of a cell line-specific gold standard interaction dataset. Moreover, classifier accuracy is influenced by the size of the feature set, with more proximity labeling experiments enhancing performance. Applying this classifier to the Jurkat surfaceome yielded over 6,000 protein associations, encompassing both known and uncharacterized interactions. As curated protein interaction databases were used for classifier training, enrichment of known interaction pairs among high scoring predictions should be interpreted as evidence that the classifier recovers features consistent with existing interaction knowledge, rather than as fully independent validation of novel association discovery. The ability of this approach to prioritize previously uncharacterized cell surface associations therefore requires orthogonal experimental or structural follow up. With this in mind, we identified and characterized a functional association between the beta subunit of the IL10 receptor (IL10RB) and a subunit of the type I interferon signaling system. Knocking out IL10RB enhanced proximal interferon signaling, as evidenced by increased STAT1 phosphorylation and elevated IRF9 levels following IFNα2 stimulation. However, this amplified signaling did not translate into improved antiviral protection. Instead, IL10RB knockout cells exhibited altered induction of 63 interferon-stimulated genes (ISGs) and increased susceptibility to vesicular stomatitis virus (VSV) infection. Decreased stimulation of the known anti-VSV factor MX1 and broadly anti-viral MX2 proteins in the IL10RB KO line likely explain this phenotype^57,58^. These findings suggest that IL10RB may influence the cellular response to type I interferon stimulation. While the increased STAT1 phosphorylation and IRF9 induction observed in the IL10RB knockout line are consistent with the broader anti-inflammatory role of IL-10/STAT3 signaling, the observation that enhanced signaling coincides with reduced antiviral protection indicates that the relationship between IL10RB and interferon signaling is more complex.^55^ At present, it remains unclear whether IL10RB directly influences the composition of the ISG response or whether these effects arise indirectly through altered receptor organization or signaling dynamics. Interestingly, the delayed induction of several ISGs observed in the IL10RB knockout cells parallels differences reported between type I and type III interferon signaling, where reduced or delayed ISG activation is associated with less potent antiviral activity^49^.

Mapping surface antigens on two cell types enabled a comparison of both antigen presence and conserved spatial arrangement across cell types. Despite limited conservation of antigens themselves, interaction networks were highly conserved. Particularly notable was the clustering of receptor ligands on Daudi cells, which included T-cell activation ligands such as ICOS ligand, LFA-3 (CD2 ligand), and TNFSF8 (CD30 ligand). This clustering may represent a regulatory mechanism whereby B cells spatially organize immune activation signals, potentially dispersing them in response to specific cues.

Finally, we leveraged the proximity labeling data to model the spatial distribution of the T cell surface proteome. This analysis revealed surface coverage patterns for antigens, distinguishing highly dispersed proteins, such as CD45, from those with more restricted distributions, such as CD226. Interestingly, our findings indicate that surface protein distribution appears largely independent of protein abundance. This observation suggests the involvement of active mechanisms in the sequestration or spatial regulation of surface proteins, rather than passive diffusion alone. These observations provide a framework for using cell surface PL to profile surface proteins that may be anchored in place, and/or confined within hypothesized cell surface pickets^56^.

It should be noted that this approach provides a protein-centric view of the cell surface without consideration of other biomolecules, such as lipids and glycans, that play critical roles in membrane structure and function. Nonetheless, given the expected density of approximately 30,000 proteins per µm^2^ on mammalian cell membranes^7^, studying the cell surface proteome alone offers a rich and informative snapshot of cell membrane topology. 3D modeling in this format makes the resolution of mapping surface distribution directly dependent upon the amount of bait proteins used for proximity labeling. Depending on the application, a 3D network with as few as 10 bait proteins might be useful for mapping and comparing cell surfaceomes. This approach is amenable to studying the cell surface interactome surrounding any given surface protein. However, when a protein is introduced or overexpressed, its membrane distribution may not faithfully reflect its native spatial organization, as elevated expression levels could lead to more diffuse localization across the plasma membrane. Additionally, the absence of endogenous interaction partners may promote non-native associations, potentially confounding interpretation of the resulting interaction network. We acknowledge that our dataset is not an exhaustive human surfaceome and that there are surface proteins that we could not detect due to limitations in the profiled cell types, the antibody/bait panel used for proximity labeling, and the sensitivity of the proteomics workflow.

Overall, analyzing surface protein distributions using large-scale PL datasets opens avenues for addressing longstanding questions in membrane biology. This methodology provides an evocative lens through which we can study how surface proteins are organized and distributed, yielding insights into the dynamic and functional architecture of the cell surface.

## Resource availability

### Lead contact

Brendan Floyd -

Carolyn Bertozzi -

### Materials availability

Knockout cell lines generated in this study may be made available upon request following institutional approval of a material transfer agreement (MTA).

## Acknowledgements

We thank Ciaran Seath (UF Scripps) for generous gifts of second and third generation Iridium photocatalysts. We also thank Nicholas Riley, Johannes Hevler, Mariko Morimoto, and David Roberts for helpful discussions.

## Author contributions statement

B.M.F. and C.R.B. conceived the project. B.M.F. and C.R.B. wrote the paper with input from E.L.S., N.A.T., J.L.Y., P.L., B.M.G., and R.A.F.. B.M.F. performed proximity labeling experiments, mass spectrometry data acquisition, immunofluorescence studies, performed data analysis, and made data visualizations. P.L. created the cellsurfacemap.org website. E.L.S. helped with proximity labeling experiments. B.M.G. and R.A.F. performed cell surface TFAM staining on peripheral blood mononuclear cells. N.A.T. synthesized and validated photocatalyzed proximity labeling materials. J.L.Y. contributed to knockout cell line generation and validation. B.S.W. and J.E.C. planned and performed viral replication assays. C.R.B supervised the project.

## Inventory of Supporting Information

Supplementary Tables S1 to S6 Movie S1

Source Data 1 – Source data for Figure 1

Source Data 2 – Source data for Figure 2

Source Data 3 – Source data for Figure 3

Source Data 4 – Source data for Figure 4

Source Data 5 – Source data for Figure 5

Source Data 6 – Source data for Figure 6

Source Data 7 – Source data for Extended Data Figure 1

Source Data 8 – Source data for Extended Data Figure 2

Source Data 9 – Source data for Extended Data Figure 3

Source Data 10 – Source data for Extended Data Figure 4

Source Data 11 – Source data for Extended Data Figure 5

Source Data 12 – Source data for Extended Data Figure 6

Source Data 13 – Uncropped image of the blot in Figure 5C

Source Data 14 – Uncropped image of the blot in Extended Data Figure 3D

## Methods

### Cell lines

The cell lines Jurkat E6-1 (ATCC #TIB-152), Daudi (ATCC #CCL-213), U-937 (ATCC CRL-1593.2), NCI-H82 (ATCC #HTB-175), K-562 (ATCC #CCL-243), Toledo (ATCC #CRL-2631), Raji (ATCC #CCL-86), COLO 205 (ATCC #CCL-222), ZR-75-1 (ATCC #CRL-1500), and A549 (ATCC #CCL-185) were cultured in T75 flasks (Genesee Scientific #25-209) in media consisting of RPMI-1640 (Thermo Scientific #11875119), 10% fetal bovine serum (FBS) (Thermo Scientific #A5256801), and 1% Penicillin-Streptomycin (Fisher Scientific #sv30010). The cell lines HeLa (ATCC #CCL-2) and HEK-293T (ATCC #CRL-3216) lines were cultured in T75 flasks in media consisting of DMEM (Thermo Scientific #11995073), 10% FBS, and 1% Penicillin-Streptomycin. The NIH:OVCAR3 (ATCC #HTB-161) cell line was cultured in a T75 flask with media consisting of RPMI-1640, 20% FBS, 1% Penicillin-Streptomycin, and 0.01 mg/mL bovine insulin (Sigma-Aldrich #I0516). All cell lines were purchased from the American Type Culture Collection (ATCC) and incubated at 37^○^C with 5% CO_2_. HeLa, U937, ZR-75-1, OVCAR-3, K-562, A549, and Raji cells are all on the International Cell Line Authentication Committee list of known misidentified cell lines. To ensure these were the expected cell lines, all these lines were acquired directly from ATCC and were used for a single experiment (Figure 3D) with an experimental outcome independent of the cells used.

### Knock-Out Cell Line Generation

The CRISPR-Cas9 mediated TFAM, IL10RB, and TRAC knockout Jurkat cell pool was generated using Synthego Gene Knockout Kit v2 (Synthego Corporation). Cells were electroporated with Streptococcus pyogenes Cas9 (Sp-Cas9, Synthego Corporation) precomplexed to sgRNAs (9:1 sgRNA:SpCas9 ratio) targeting TFAM, IL10RB, and TRAC in the SE buffer (Lonza #V4XC-1032) using the Lonza 4D Nucleofector X on the CL-120 setting following Synthego’s instructions. Cells were expanded for one week before and for the IL10RB knockout line, IL10RB-positive cells were removed by FACS following staining by anti-Hu IL10RB (S17009F, 1 μg/100 μL, Biolegend #396802) with secondary anti-mouse IgG Alexa Fluor 647 (2 μg/mL, Thermo Scientific #A-21235). sgRNA guides: IL10RB 5′-GGUACUGAGCUGUGAAAGUC-3′; 5′-UAUUUUCAUAGCAUUGGGAA-3′; 5′-AUUUCAAGAACAUUCUACAG-3′. TRAC 5’-CUCUCAGCUGGUACACGGCA-3’, 5’-GAGAAUCAAAAUCGGUGAAU-3’, 5’-ACAAAACUGUGCUAGACAUG-3’. TFAM 5’-UGGCGUUUCUCCGAAGCAUG-3’; 5’-GGCAAGCGGGCCUACCUGAA-3’; 5’-UAGGGUCUACAUUCCAACCC-3’. Knockout cell lines generated in this study were kept at -80^○^C in Bambanker Cell Freezing Media (Bulldog Bio #BB01) and may be made available upon request following institutional approval of a material transfer agreement (MTA).

### Flow cytometry cell surface staining

Cells were cultured in media overnight (37^○^C, 5% CO_2_) in a T75 flask. Cells were moved to 15 mL conical tube and were counted using Trypan Blue (VWR #VWRVK940) and a Countess Cell Counter (Thermo Scientific). Cells were then diluted to a concentration of 5 million cells/mL in FACS buffer (PBS pH 7.4 and 0.1% w/v bovine serum albumin). 200 μL of the cell solution (or 1 million cells) was then transferred to a 96-well V bottom plate (Fisher Scientific #0720096). Cells were kept on ice and spun at 4^○^C for the remainder of the protocol. Cells were then spun at 300xg for 3 min. Supernatant was aspirated and cells were washed 2X with 200 μL FACS buffer (0.1% BSA in PBS). Cells were pelleted by spinning at 300xg for 3 min after each wash. After aspirating following the second wash, cells were incubated in 100 μL of FACS buffer containing the primary antibody diluted to 0.01 mg/mL. Cells were then incubated on ice for 30 min covered from light. Cells were then pelleted by spinning at 300xg for 3 min. The supernatant was then aspirated and the cells were then washed twice with 200 μL FACS buffer (0.1% BSA in PBS). After aspirating following the second wash, cells were incubated in 100 μL of FACS buffer containing the secondary antibody diluted to 2 μg/mL. Cells were then incubated on ice for 30 min covered from light. Cells were then pelleted by spinning at 300xg for 3 min. The supernatant was then aspirated and cells were washed twice with 200 μL FACS buffer (0.1% BSA in PBS). After aspirating, following the second wash cells were incubated in 100 μL of FACS buffer containing Sytox red (Thermo Scientific #S34859) diluted by 1:1000 v/v to a final concentration of 4 nM. Cells were then pelleted by spinning at 300xg for 3 min. The supernatant was then aspirated and cells were resuspended in 100 μL of FACS buffer. For adherent cell lines, cells were resuspended in 100 μL of FACs buffer containing 1 mM EDTA (Thermo Scientific #15575020). Fluorescence was then monitored using a MACSQuant Analyzer 10 (Miltenyi Biotec) and analysis was done using the FlowJo software package. Gating was performed on single cells and live cells.

### Donor PBMC preparation and cell surface staining

A leukocyte filtration device was cut on both ends to allow drainage of blood. 30 mL of PBS and 20 mL of dead volume were pipetted through the distal end of the filters to fully eject leukocyte rich blood. This mixture of PBS and blood was spun at 500xg for 10 min, yielding a buffy coat. The buffy coat was spun again to further remove any red blood cells. Buffy coats were resuspended in FACS buffer (PBS pH 7.4 and 0.1% w/v bovine serum albumin). Cells were incubated with Human TruStain FcX (BioLegend, 422302) blocking solution; 5μL of blocking solution was added for every 1e6 cells in 100μL. Cells were then stained with 500 ng of rabbit IgG isotype control (Novus Biologicals #NBP2-24893) or anti-TFAM (Proteintech #23996-1-AP) antibodies in 100 μL. Cells were incubated for 30 min on ice. They were washed once with FACS buffer and then stained with goat anti-rabbit IgG Alexa Fluor 647 and incubated for 30 min on ice. They were then washed once with FACS buffer and then stained with lineage antibodies CD3 FITC for T-cells (BioLegend #300452), CD19 PE-Cy7 for B-cells (BioLegend #302216), CD14 PerCP-Cy5.5 for monocytes (BioLegend #325622), and CD66b BV421 for neutrophils (BioLegend #392916) at a concentration of 1:100. Lineage antibody staining was performed on ice for 30 min. Cells were washed twice with FACS buffer and resuspended in DAPI containing buffer just prior to flow cytometry (BD LSRFortessa). Cells were gated first to yield singlets and then subsequently DAPI negative cells were gated to analyze total live cells. Afterwards, CD3 and CD19 single positive cells were gated; the double negative fraction was analyzed to look at CD14 and CD66b positive cells.

### Interferon stimulation and reading out IRF9 abundance using intracellular flow cytometry

Jurkat E6-1 cells were cultured overnight at (37^○^C 5% CO_2_) in a T75 flask. Cells were then transferred to 1.5 mL microcentrifuge tubes and pelleted at 300xg for 5 min and resuspended in media at 500,000 cells/mL. 500,000 cells (or 1 mL of the resuspended solution) were then added to a 6 well plate (Corning #3516). IFNα2 (Thermo Scientific #300-02AA-20UG) was then added at the specified concentration from a stock concentration of 1 μg/mL in PBS pH 7.4. For the blank negative control 10 μL of sterile PBS pH 7.4 was added to the media. Cells were then incubated overnight at 37^○^C with 5% CO_2_. Cells were then transferred to 1.5 mL microcentrifuge tubes and pelleted at 300xg for 5 min. Cells were then counted and resuspended in FACs buffer to make a 5 million cell/mL solution. 200 μL of the cell solution (or 1 million cells) were then transferred to a 96-well V bottom plate. Cells were then spun at 300xg for 3 min. The supernatant was then aspirated and cells were resuspended in 200 μL of PBS. Cells were then spun at 300xg for 3 min. The supernatant was then aspirated and cells were resuspended in 100 μL of Zombie Violet (Biolegend #423113) diluted in PBS by 1:1,000. Resuspended cells were then incubated in the dark for 20 min at 22^○^C. Cells were then pelleted by spinning at 300xg for 3 min and the supernatant was aspirated. Cells were resuspended in 200 μL of FACS buffer and spun at 300xg for 3 min. The supernatant was then aspirated and cells were fixed by adding 200 μL Fixation Buffer (Biolegend #426803) and incubated in the dark for 20 min at 22^○^C. Cells were then spun at 300 xg for 5 min and the supernatant was aspirated. 10X Intracellular Staining Perm Wash Buffer (Biolegend #426803) was diluted to 1X in Deionized water. The fixed cells were then resuspended in 200 μL of the diluted Intracellular Staining Perm Wash Buffer and centrifuged at 300xg for 5 min. This washing step was repeated two more times. Anti-IRF9 mouse monoclonal primary antibody (Biolegend #660702) was then diluted to a final concentration of 0.01 mg/mL in 1X Intracellular Staining Perm Wash Buffer. After the final wash, cells were resuspended in the diluted antibody solution and incubated for 30 min at 22^○^C. Cells were then washed twice with 200 μL of 1X Intracellular Staining Perm Wash Buffer and centrifuged at 300xg for 5 min and the supernatant was aspirated between each wash. Goat anti-mouse IgG Alexa Fluor 488 secondary antibody (Thermo Scientific #A32723) was then diluted to a final concentration of 2 μg/mL in 1X Intracellular Staining Perm Wash Buffer. After the final wash cells were resuspended in the diluted secondary antibody solution and incubated for 30 min at 22^○^C. Cells were then washed twice with 200 μL of 1X Intracellular Staining Perm Wash Buffer and centrifuged at 300xg for 5 min between each wash. Cells were then resuspended in 100 μL FACs buffer for flow cytometry analysis. Fluorescence was then monitored using a MACSQuant Analyzer 10 (Miltenyi Biotec) and analysis was done using the FlowJo software package. Gating was performed on single cells and live cells.

### Interferon stimulation and reading out phosphorylated STAT1 abundance using Western blot

Jurkat E6-1 cells were cultured overnight at (37^○^C 5% CO_2_) in a T75 flask. Cells were then transferred to 1.5 mL microcentrifuge tubes and pelleted at 400xg for 4 min and resuspended in media at 2 million cells/mL. IFNα2 (Thermo Scientific #300-02AA-20UG) was then added at 1 or 10 ng/mL from a stock concentration of 1 μg/mL in PBS pH 7.4. For the blank negative control 10 μL of sterile PBS pH 7.4 was added to the media. Cells were then incubated for 30 min at 37^○^C on a Thermomixer (Fisher Scientific #05-412-503) shaking at 0.15xg. Cells were then transferred to 1.5 mL microcentrifuge tubes and pelleted at 300xg for 5 min. Cells were then pelleted at 400xg for 4 min and the supernatant was aspirated. Cells were then resuspended in 1 mL of PBS and the cells were pelleted at 400xg for 4 min. The supernatant was aspirated and cells were resuspended in 100 µL RIPA buffer containing 1X protease inhibitor and 1X phosphatase inhibitor. Cells were lysed at 4^○^C for 20 min. The samples were then spun at 16,000xg for 15 min at 4°C and the supernatant was moved to a new LoBind tube. The protein concentration was measured by BCA (Pierce BCA Assay, Thermo Scientific #23225). 10 µg of protein in lysate was denatured with a 4X loading buffer which was supplemented with 10% v/v 1 M dithiothreitol (Thermo Scientific #15-508-013) and boiled at 95 °C for 10 min. 20 µL of the reduced protein sample was then loaded on a 12-well SDS–PAGE gel (10% Bis-Tris gel, Bio-Rad #345-0123) and run at 200 V for 60 min in 1X XT MOPS buffer (Bio-Rad #1610788). Protein was then transferred from the gel to a 0.45 µm nitrocellulose membrane (Bio-Rad #1620115). After transfer, total protein was measured by staining with total protein stain (Revert 700 Total Protein Stain, LI-COR Biosciences #926-11021) and incubated for 3 min at 22°C. The membrane was then imaged using an OdysseyCLxImager (LI-COR Biosciences). The total protein stain was then removed using a basic buffer and washed three times with PBS pH 7.4. The membrane was then blocked with PBS Odyssey Blocking Buffer (LI-COR Biosciences #927-40000) for 1 h at room temperature with gentle shaking. The membrane was stained with anti-pSTAT1 (BioLegend #666402) diluted by 1:500 to a final concentration of 1 µg/mL in PBS Odyssey Blocking Buffer overnight at 4°C with gentle shaking, then washed four times with PBS with 0.1% Tween-20 (Bio-Rad #1706531), PBS-T, for 5 min each. The membrane was then incubated with 800CW goat anti-mouse IgG (LI-COR Biosciences #926-32210) diluted by 1:1,000 to a concentration of 1 µg/mL in PBS Odyssey Blocking Buffer for 1 h at room temperature with gentle shaking. The membrane was washed four times with PBS-T and then imaged using an OdysseyCLxImager (LI-COR Biosciences).

### Surface antigen staining for confocal microscopy

Jurkat E6-1 cells were cultured overnight (37C, 5% CO2) in a T75 flask. Cells were moved to 15 mL conical tube and were counted using Trypan Blue (VWR #VWRVK940) and a Countess Cell Counter (Thermo Scientific). Cells were then diluted to a concentration of 5 million cells/mL in FACS buffer (PBS pH 7.4 and 0.1% w/v bovine serum albumin). 200 µL of the cell solution, or 1 million cells, were then transferred to a 96-well V bottom plate. Cells were kept on ice and spun at 4^○^C for the duration of the protocol. Cells were then spun at 300xg for 3 min. The supernatant was then aspirated and cells were washed two times with 200 µL FACS buffer (0.1% BSA in PBS). Cells were pelleted by spinning at 300xg for 3 min after each wash. After aspirating the supernatant following the second wash, cells were incubated in 100 µL of FACS buffer containing the primary antibody diluted to 0.01 mg/mL. Cells were then incubated on ice for 30 min covered from light. Cells were then pelleted by spinning at 300xg for 3 min. The supernatant was then aspirated and cells were washed two times with 200 µL of FACS buffer. After aspirating following the second wash, cells were incubated in 100 µL of FACS buffer containing the secondary antibody goat anti-mouse IgG Alexa Fluor 488 (Thermo Scientific #A32723) diluted to 0.01 mg/mL. Cells incubated on ice for 30 min covered from light. Cells were then pelleted by spinning at 300xg for 3 min. The supernatant was then aspirated and cells were washed two times with 200 µL of FACS buffer. After aspirating following the second wash, cells were incubated in 100 µL of FACS buffer containing diluted CellMask Orange Plasma Membrane Stain (Thermo Scientific #C10045) diluted by 1:10000 to a final concentration of 0.5 µg/mL in FACS and incubated on ice in the dark for 5 min. Cells were then pelleted by spinning at 300xg for 3 min. The supernatant was aspirated and the cells were washed two times with 200 µL FACS buffer. For fixation, cells were resuspended in a solution of 4% v/v paraformaldehyde in PBS and incubated in the dark for 30 min at room temperature. Cells were then pelleted by spinning at 300xg for 3 min. Cells were then washed one time with 200 µL of PBS and pelleted again and the supernatant was aspirated. Cells were then resuspended in a solution of DAPI (Fisher Scientific #EN62248) diluted by 1:1000 to a final concentration of 5 µg/mL in PBS and incubated in the dark for 5 min at room temperature. Cells were then pelleted and resuspended in 25 µL of PBS. 25 µL of the cell solution was then pipetted onto a glass slide and 25 µL of mounting medium (Fisher Scientific #NC1601055) was then mixed with the cells. A coverslip was then placed over the solution and the cells were enclosed using clear nail polish (Electron Microscopy Sciences #72180). The slides were then left to dry for 16 h in the dark at room temperature.

### TFAM-mClover plasmid lentivirus packaging

The plasmid pcDNA3 TFAM-mClover was obtained from Addgene (Addgene #129574). 2-3 days prior to packaging, Lenti-x 293T cells were cultured on a 10 cm plate in DMEM media with 10% fetal bovine serum (FBS) and incubated at 37^○^C with 5% CO_2_. Cells were plated 24 h prior to transfection and were grown to at least 80% confluency. For lentiviral packaging, Cellecta’s ready-to-use lentiviral packaging plasmid mix (Cellecta #CPCP-K2A) and 2 ug of the TFAM-mClover plasmid were mixed together in a sterile 2 mL polypropylene tube. DMEM media was then added to the plasmid mixture. PLUS reagent was added, mixed, and incubated at room temperature for 15 min. 30 uL of Lipofectamine 3000 Transfection reagent (Thermo Scientific #L3000001) was diluted in 1 mL of DMEM to make a master mix and the diluted lipofectamine was then added to the DNA/PLUS reagent complex and mixed gently. The solution was incubated at room temperature for 15 min. 2.5 mL of the mixture was then added to each flask with care taken not to dislodge the cells from the flask. Cells were incubated for 24 h at 37^○^C with 5% CO_2_. After the 24 h incubation, DMEM media was replaced with fresh DMEM supplemented with 10% FBS,

### TFAM-mClover HEK-293T transduction

Wild type HEK-293T cells were diluted to 2 million cells/mL and were plated on a 12-well plate. 20 mM HEPES was added for pH balance. Polybrene was added to a concentration of 8 ug/mL from an initial stock of 10 mg/mL. Lentivirus containing the TFAM-mClover plasmid was added to the plate. The plate was parafilmed and spun at 1200 xg at 25^○^C for 2 h. The parafilm was removed and the plate was incubated overnight at 37^○^C with 5% CO_2_. 500 uL of media was added 3 h after incubation was started. The next day, cells were pelleted and resuspended in fresh media. After cells reached full confluency, cells were sorted at the Stanford FACS core facility. HEK-293T TFAM-mClover cells generated in this study were kept at -80^○^C in Bambanker Cell Freezing Media (Bulldog Bio #BB01) and may be made available upon request following institutional approval of a material transfer agreement (MTA).

### TFAM-mClover HEK-293T confocal staining

HEK 293T cells were plated 18-24 h before treatment into 8-well labtek plates with a confluency of approximately 30,000 cells/well. After cells reached desired confluency, media was aspirated from each of the wells and a prewarmed (37^○^C) MitoTracker Red (Cell Signaling #8778S) diluted in DMEM to 100 nm was added to the cells and cells were incubated for 30 min at 37^○^C. Following incubation, cells were washed once with PBS and then 100 ul of CellMask Orange plasma membrane stain (Thermo Scientific #C10045) diluted at 1:30,000 in PBS was added. This was incubated for 5 min at 37^○^C. Cells were then washed twice in PBS. Following this, cells were incubated with 1 ug/mL Hoescht at a 1:1,000 dilution in PBS for 15 min. The Hoescht solution was removed after the incubation, and cells were washed once with PBS. Cells were then left in PBS for confocal imaging.

### Confocal microscopy

A Nikon A1R confocal microscope equipped with a Pan Fluor 60x oil immersion 1.30-numerical aperture objective (Nikon) was used. This instrument is equipped with a 405-nm violet laser, a 488-nm blue laser, a 561-nm green laser and a 639-nm red laser.

### Wheat germ agglutinin-horseradish peroxidase (WGA-HRP) catalyzed cell surface biotinylation

Jurkat E6-1 were used at 10 million cells/labeling experiment and, unless otherwise noted, were pelleted at 1,000xg for 4 min at 4 °C. Cells were first removed from culture, washed twice with cold DPBS (Fisher Scientific #MT21031CV) and resuspended in cold DPBS at 10 million cells/mL and transferred in 1 mL aliquots to Protein LoBind tubes (Fisher Scientific #13-698-794). The cells were then pelleted to remove the supernatant and resuspended in 1 mL of cold DPBS containing 5 µg of WGA-HRP (Vector Laboratories #PL-1026-2) or 5 µg of unconjugated HRP (Thermo Fisher Scientific #31490). The cells were then incubated at 4°C for 1 h on a rotisserie, followed by centrifugation to remove the supernatant and then washed twice in cold DPBS. Each sample was resuspended in 1 mL of cold DPBS and added directly to a 15 mL conical tube containing 4 mL of reaction buffer,250 µM biotinyl tyramide (Sigma-Aldrich #SML2135-250MG) and 1 mM H_2_O_2_ (Sigma-Aldrich #H1009-100ML) in cold DPBS, and incubated for 1 min at 4 °C. After 30 seconds, 5 mL quenching buffer, DPBS containing 5 mM Trolox (Sigma-Aldrich #238813-5G), 10 mM Sodium L-ascorbate (Sigma-Aldrich #11140-250G), 10 mM NaN_3_ (Sigma-Aldrich #S2002-100G), was added to the cells. The cells were then pelleted by centrifugation (4 min at 1,000xg, 4°C) and washed once with 10 mL of quenching buffer. After the final wash, the cells were pelleted and each sample was resuspended in 1 mL of RIPA buffer (Pierce RIPA Buffer, Thermo Scientific #89901) containing 1X protease inhibitors (cOmplete Protease Inhibitor Cocktail EDTA-Free, Roche #04693159001) and incubated for 20 min at 4 °C. The samples were then spun at 16,000xg for 15 min at 4 °C. The supernatant was then moved to a new tube and the protein concentration was measured by BCA (Pierce BCA Assay, Thermo Scientific #23225) and samples were stored at -80°C until streptavidin bead enrichment.

### Horseradish peroxidase catalyzed cell surface biotinylation

Jurkat E6-1 or Daudi cells were used at 50 million cells/labeling experiment and, unless otherwise noted, were pelleted at 1,000xg for 4 min at 4 °C. Cells were first removed from culture, washed twice with cold DPBS (Fisher Scientific #MT21031CV) and resuspended in cold DPBS at 50 million cells/mL and transferred in 1 mL aliquots to Protein LoBind tubes (Fisher Scientific #13-698-794). The cells were then pelleted to remove the supernatant and resuspended in 1 mL of cold DPBS containing 25 µg of isotype control (Thermo Scientific #31903) or 25 µg of the respective targeting antibody. The cells were then incubated at 4°C for 30 min on a rotisserie, followed by centrifugation to remove the supernatant and then washed twice in cold DPBS. The cells were then resuspended in 1 mL of cold DPBS containing 25 µg goat α-Mouse IgG HRP (EMD Millipore #12-349) and incubated for 30 min at 4°C on a rotisserie. The cells were then pelleted to remove the supernatant and washed twice in cold DPBS. Each sample was resuspended in 1 mL of cold DPBS and added directly to a 15 mL conical tube containing 4 mL of reaction buffer,250 µM biotinyl tyramide (Sigma-Aldrich #SML2135-250MG) and 1 mM H_2_O_2_ (Sigma-Aldrich #H1009-100ML) in cold DPBS, and incubated for 1 min at 4 °C. After 1 min, 5 mL quenching buffer, DPBS containing 5 mM Trolox (Sigma-Aldrich #238813-5G), 10 mM Sodium L-ascorbate (Sigma-Aldrich #11140-250G), 10 mM NaN_3_ (Sigma-Aldrich #S2002-100G), was added to the cells. The cells were then pelleted by centrifugation (4 min at 1,000xg, 4°C) and washed once with 10 mL of quenching buffer.

After the final wash, the cells were pelleted and each sample was resuspended in 1 mL of membrane permeabilization buffer (MEM-PER Plus Membrane Fractionation Kit, Thermo Scientific #89842) containing 1X protease inhibitors (cOmplete Protease Inhibitor Cocktail EDTA-Free, Roche #04693159001) and incubated for 20 min at 4 °C. The samples were then spun at 16,000xg for 15 min at 4 °C. The supernatant enriched in cytosolic proteins was removed, and the pellet was resuspended in 300 µL lysis buffer (Pierce RIPA Buffer, Thermo Scientific #89901) containing 1% SDS v/v and 1X protease inhibitors. The samples were sonicated in the lysis buffer to break up the membrane pellet once for 5 s at power level 6 using a probe sonicator and then heated for 5 min at 95 °C. The samples were then diluted to 1 mL with RIPA and sonicated twice for 5 s at power level 3 using a Misonix Sonicator 3000 probe sonicator. The protein concentration was measured by BCA (Pierce BCA Assay, Thermo Scientific #23225) and samples were stored at -80°C until streptavidin bead enrichment.

### Iridium catalyzed cell surface biotinylation (μMap)

Jurkat E6-1 cells were used at 50 million cells/labeling experiment and, unless otherwise noted, were pelleted at 1,000xg for 4 min at 4 °C. Cells were first removed from culture, washed twice with cold DPBS (Fisher Scientific #MT21031CV) and resuspended in cold DPBS at 50 million cells/mL and transferred in 1 mL aliquots to Protein LoBind tubes (Fisher Scientific #13-698-794). The cells were then pelleted to remove the supernatant and resuspended in 1 mL of cold DPBS containing 25 µg of isotype control (Thermo Scientific #31903) or 25 µg of the respective targeting antibody. The cells were then incubated at 4°C for 30 min on a rotisserie, followed by centrifugation to remove the supernatant and then washed twice in cold DPBS. The cells were then resuspended in 1 mL of cold DPBS containing 25 µg of iridium-conjugated goat α-mouse secondary IgG (Supplemental Methods). The samples were incubated on a rotisserie for 30 min at 4 °C and then pelleted to remove the supernatant. The cells were washed twice with 1 mL of cold DPBS and then resuspended in 1 mL of cold DPBS containing 250 µM diazirine biotin (Supplemental Methods). The samples were placed on the Lumidox II photoreactor (Analytical #LUM2CON) and irradiated on a 24-well LED array (Analytical #LUM22418LA445) with 445 nm light at 195W for 10 min at 4 °. After irradiation, the cells were then pelleted to remove the supernatant and washed twice with 1 mL of cold DPBS.

After the final wash, the cells were pelleted and each sample was resuspended in 1 mL of membrane permeabilization buffer (MEM-PER Plus Membrane Fractionation Kit, Thermo Fisher Scientific #89842) containing 1X protease inhibitors (cOmplete Protease Inhibitor Cocktail EDTA-Free, Roche #04693159001) and incubated for 20 min at 4 °C. The samples were then spun at 16,000xg for 15 min at 4 °C. The supernatant enriched in cytosolic proteins was removed and the pellet was resuspended in 300 µL of lysis buffer (Pierce RIPA buffer, Thermo Scientific #89901) containing 1% v/v SDS and 1X protease inhibitors. The samples were sonicated in the lysis buffer to break up the membrane pellet once for 5s at power level 6 using a Misonix Sonicator 3000 probe sonicator and then heated for 5 min at 95°C. The samples were then diluted to 1 mL with RIPA and sonicated twice for 5 s at power level 3 using a Misonix Sonicator 3000 probe sonicator. The protein concentration was measured by BCA and samples stored at -80°C until streptavidin bead enrichment.

### Magnetic streptavidin bead enrichment

For bead enrichment membrane fractions were added to a Protein LoBind tube containing 125 µL of streptavidin magnetic beads (Thermo Fisher Scientific #88817) that were pre-washed twice with 500 µL of RIPA buffer. The samples were then incubated for 3 h at room temperature on a rotisserie and the beads were then pelleted on a magnetic rack (Thermo Scientific #12321D). The supernatant was removed, and the beads were washed three times with 500 µL 1% v/v SDS (Thermo Scientific #15553027) in DPBS, three times with 500 µL 1M NaCl (Fisher Scientific #S6713) in DPBS, three times with 500 µL 10% v/v EtOH (Gold Shield Distributors #412804) in DPBS, and once with 500 μL RIPA buffer (Thermo Scientific #89901). The beads were incubated in each of the washes for 3 min before pelleting to remove the wash. After the final wash, the beads were resuspended in 25 µL of 5% v/v SDS containing 20 mM DTT and 25 mM biotin (Sigma-Aldrich #B4501-1G) and heated to 95°C for 10 min. The samples were placed on the magnetic rack and the supernatant was collected and transferred to a new Protein LoBind tube and stored at -80 °C until quantitative proteomic sample preparation and analysis

### Tandem mass spectrometry sample preparation

75 μg of biotin-enriched membrane samples were reduced in a solution of 5% v/v SDS (Thermo Fisher Scientific #15553027) with 10 mM dithiothreitol (Bio-Rad #1610611) and incubated at 95°C for 10 min. Samples were then allowed to cool to room temperature and then alkylated using 40 mM 2-chloroacetamide (Sigma-Aldrich #C0267-500G) and incubated for 1 h at room temperature. The alkylation reaction was quenched by adding 85% (m/v) phosphoric acid (Millipore Sigma #W290017-1KG-K) to a final solution percentage of 1.2% phosphoric acid in the alkylated protein solution. 700 μL (7X volume) of S-trap bind wash buffer, 100 mM triethylammonium bicarbonate buffer (TEAB) (Fisher Scientific #60044974), 90% (v/v) MeOH (Fisher Scientific #A452-4), was then added to the alkylated sample. Samples were then loaded onto a micro-S-trap-column (Protifi #C02-micro-80) in 175 μL increments and spun at 4000xg for 20 s for each loading cycle. Sample caught on the column was then washed three times using the S-trap bind wash buffer with spins of 4000xg for 30 s to pass the buffer over the column. Mass spectrometry-grade trypsin (Promega #V5113) was then diluted in 50 mM TEAB buffer to a concentration of 0.15 mg/mL. 25 μL of the diluted trypsin solution, or 3.75 μg of trypsin, was loaded onto each column to make a final trypsin:protein ratio of 1:20. Samples were then incubated at 47°C for 90 min. The column was then washed using 50 mM TEAB followed by 0.2% (v/v) formic acid (Thermo Fisher #28905) in water with a spin at 1000xg for 60 s between each wash. Digested samples were then eluted by adding 40 μL of a solution of 50% v/v acetonitrile (Fisher Scientific #A955-212), 0.2% v/v formic acid (Thermo Fisher #28905), and 50% v/v water then spun at 4000xg for 60 s. Samples were then dried using vacuum centrifugation (CentriVap Complete Vacuum Concentrator, Labconco #7315023) for 16 h. The samples were resuspended in 0.2% (v/v) formic acid diluted in water immediately prior to mass spectrometry data acquisition.

### Tandem mass spectrometry data acquisition

All samples were resuspended in a 0.2% (v/v) formic acid diluted in water at 1 μg/μl, and 1 μg of each sample was added to a pooled sample tube for the six gas phase fractionation injections. 1 μL (1 μg) of the total peptide solution was injected in the column for each sample. Peptides were separated over a 25-cm EasySpray reversed phase LC column (75 μm inner diameter packed with 2 μm, 100 Å, PepMap C18 particles, Thermo Fisher Scientific #ES802). The mobile phases (A: water with 0.2% (v/v) formic acid and B: acetonitrile with 0.2% (v/v formic acid) were driven and controlled by a Dionex Ultimate 3000 RPLC nano system (Thermo Fisher Scientific). Gradient elution was performed at 300 nL/min. Mobile phase B was increased from 1 to 5% (v/v) over 6 min, followed by a gradual increase to 25% by 60 min, and ending with a ramp to 90% B at 71 min, and a wash at 90% B for 5 min. Flow was then ramped back to 1% B over the course of 1 min, and the column was re-equilibrated at 1% B for 15 min, for a total analysis of 90 min. Eluted peptides were analyzed on an Orbitrap Fusion Tribrid MS system (Thermo Fisher Scientific). Precursors were ionized using an EASY-Spray ionization source (Thermo Fisher Scientific) held at +2.2 kV compared to ground, and the column was held at 40 °C. The inlet capillary temperature was held at 275 °C. For gas phase fractionation injections, a pooled sample was injected six times and acquired using staggered-window injections with m/z windows of 395 to 505 m/z, 495 to 605 m/z, 595 to 705 m/z, 695 to 805 m/z, 795 to 905 m/z, and 895 to 1005 m/z. For all staggered-window injections a 4 m/z precursor isolation window was used. MS/MS scans were collected using high-energy collisional dissociation at 33 normalized collision energy and mass analysis was performed in the Orbitrap using a resolution of 30,000 while scanning from 120–2000 m/z. Individual samples were collected using single-injection data independent acquisitions (DIA) with survey scans of peptide precursors collected in the Orbitrap from 385-1015 m/z with an automatic gain control target of 400,000, a maximum injection time of 55 ms and a resolution of 60,000. An isolation window of 16 m/z was used to select precursor ions with the quadrupole. MS/MS scans were collected using high-energy collisional dissociation at 33 normalized collision energy and mass analysis was performed in the Orbitrap using a resolution of 30,000 while scanning from 120–2000 m/z.

### Mass spectrometry database search and quantitation

Proteins were identified with DIA-NN^60^ and MSFragger 4.0^61^ software on the Fragpipe GUI version 21.0^62^, searching against the UniProt human reviewed proteome in addition to common contaminants.

Methionine oxidation (+15.995 Da) and N-terminal acetylation (+42.011 Da) were included as variable modifications, while carbamidomethylation of cysteine (+57.021 Da) was included as a fixed modification. Precursor ion search tolerance was set to 20 ppm and fragment ion mass tolerance was set to 20 ppm. Peptide and protein identifications were thresholded at a 1% false discovery rate using a target-decoy method. Proteins were quantified and normalized by label-free quantitation using the DIA-NN software. For quantitative comparisons, protein intensity values were log transformed, and missing values for proteins were imputed from a normal distribution with a width of 0.3 and a down shift value of 1.8 in Perseus (version 2.0)^63^. A two-sided t-test was additionally done using Perseus with false discovery rate (FDR) correction performed using permutation-based FDR with 250 randomizations.

### Pairwise protein subunit distance measurements

T-cell receptor subunit distances were measured in a pairwise manner across all TCR subunits from the center mass of each subunit from the cryo-EM structure of the TCR-CD3 complex (PDB 6JXR). For subunits with a stoichiometry greater than 1 (eg. 2 copies) the mean distance between protein pairs was taken.

### Unsupervised clustering and functional domain analysis

A matrix of fold change values was generated including data from our 65 proximity labeling experiments in Jurkat cells and 20 proximity labeling experiments in Daudi cells. Fold change matrices were hierarchically clustered and the resulting dendrogram was cut at intervals to obtain a nested hierarchy of associations. Cuts closer to the root of the dendrogram result in larger groups of proteins and cuts closer to the tips define smaller groups. In both Jurkat and Daudi cell lines known protein assemblies cluster together at the smallest group. In Jurkat this is represented by the CD3-TCR complex and in Daudi cells the MHC class II HLA-DR complex.

### Feature assembly for identifying surface protein associations

A matrix of fold change values containing the enrichment values for 65 proximity labeling experiments on Jurkat cells was created and filtered to contain only proteins quantified in at least 20 proximity labeling experiments. Fold changes were then normalized to lie between 0 and 1 by min-max normalization. The difference between fold change values for a given protein was then estimated on an all-by-all basis for each of 60,031 surface protein pairs. A matrix of similarity measures was then made from our fold change matrix by doing all-by-all pairwise estimations of Pearson correlation, Spearman correlation, Euclidean distance, Bray-Curtis dissimilarity, Manhattan distance, and Chebyshev distance for each surface protein pair. Euclidean distance, Bray Curtis score, Manhattan distance, and Chebyshev distance were inverted and normalized to fall between 0 and 1 using min-max normalization. The matrix of fold change differences and similarity measures were then concatenated to give a final feature matrix containing 4,202,170 data points.

### Gold standard protein interaction training and test set assembly

Due to the limited size of datasets containing known human surface protein interactions we utilized several sources to construct our gold standard interaction set: CORUM mammalian complex dataset^64^, HuMap 2.0^65^, STRING.db^66^ with interactions filtered to score above 800, and BioGRID^67^. For protein complex datasets, all complex members were converted to binary interactions where each subunit is considered an interactor to all other members of the complex. The compilation of these datasets led to a final binary gold standard interaction list of 1,185,470 protein-protein interactions Of these interactions, 1,532 comprised pairs in which both proteins were detected in the Jurkat cell surface proteome. We further filtered these pairs to retain interactions with Pearson correlation values greater than 0.3, resulting in 1,009 positive protein-pair labels used for classifier training and stratified 5-fold cross-validation.

We additionally generated a more conservative gold-standard interaction set incorporating UniProt-derived protein-protein interactions and a more stringent BioGRID multi-validation (BioGRID-MV) subset. UniProt PPIs were extracted from UniProt interaction annotations and converted to binary gene-pair interactions. For the BioGRID-MV subset, BioGRID interactions were restricted to physical interaction records and only pairs supported by two or more BioGRID records were retained. This revised gold-standard set contained 604 positive protein pairs within the Jurkat cell surface proteome after applying the same Pearson correlation filter.

### Random forest classifier design and training

For the binary prediction of protein associations, we used the scikit-learn^68^ Random Forest Classifier. We then searched for optimal hyperparameters for the classifier with 5-fold cross-validation on the labeled protein-pair feature matrix. The final random forest classifier was trained using balanced class weights, no bootstrapping, entropy-based splitting, square-root feature sampling at each split, no maximum tree depth, a minimum of 2 samples per leaf, a minimum of 5 samples required to split an internal node, and 200 estimators. The classifier was evaluated using the same stratified 5-fold cross-validation framework, where each fold was held out from model fitting and used for performance assessment. Across folds, the classifier achieved an average area under the receiver operating characteristic curve of 0.92 and an area under the precision-recall curve of 0.62. True positive rate, false positive rate, precision, recall, and false discovery rate were calculated from the cross-validated predictions. This model was applied to the entire feature matrix to give an association score for each surface protein pair with higher scores corresponding to higher likelihood of the proteins interacting or colocalizing. Because curated protein interaction databases were used to define positive training labels, enrichment of known interaction pairs among high scoring predictions should be interpreted as evidence that the classifier recovers features consistent with existing interaction knowledge, rather than as fully independent validation of novel association discovery. The ability of this approach to prioritize previously uncharacterized cell surface associations therefore requires orthogonal experimental or structural follow up.

### Classifier architecture comparison for cell surface protein association prediction

Random forest classifier performance was compared against eight additional classifier architectures: ExtraTrees, logistic regression, XGBoost, gradient boosting, histogram gradient boosting, radial basis function support vector machine, linear support vector machine, and AdaBoost. All classifiers were evaluated using the same labeled protein-pair feature matrix and identical stratified 5-fold cross-validation splits. For each model, out-of-fold prediction scores were pooled across the five folds and used to calculate precision-recall and receiver operating characteristic curves.

The random forest classifier was trained using balanced class weights, no bootstrapping, entropy-based splitting, square-root feature sampling at each split, no maximum tree depth, a minimum of 2 samples per leaf, a minimum of 5 samples required to split an internal node, and 200 estimators. ExtraTrees was trained using balanced class weights, entropy-based splitting, square-root feature sampling, no maximum tree depth, a minimum of 2 samples per leaf, a minimum of 5 samples required to split an internal node, and 300 estimators. Logistic regression was trained using L2 regularization, balanced class weights, a regularization strength of C = 1.0, the liblinear solver, and a maximum of 5,000 iterations.

Gradient boosting was trained with 200 estimators, a learning rate of 0.05, and a maximum tree depth of 3. Histogram gradient boosting was trained with 300 iterations, a learning rate of 0.05, and a maximum of 31 leaf nodes. The RBF and linear support vector machine classifiers were trained using balanced class weights and probability estimation. AdaBoost was trained with 200 estimators and a learning rate of 0.05. XGBoost was trained with 300 estimators, a learning rate of 0.05, a maximum tree depth of 3, subsampling of 0.8, column sampling by tree of 0.8, a binary logistic objective, log-loss evaluation metric, and class weighting based on the ratio of negative to positive training examples.

### AlphaFold2-Multimer heterodimer prediction

Protein pairs with random forest association scores were filtered to pairs in which both proteins had an annotated type I transmembrane domain. This left 6,903 pairs from the original 60,031 pairs. UniProt topology annotations were used to define extracellular domain (ECD) boundaries, and ECD sequences were extracted for proteins with extracellular domains of at least 30 amino acids. Pair sets were then generated from the top 100 highest-scoring random forest predictions, the bottom 100 lowest-scoring predictions, and 100 randomly sampled pairs. For each pair, the two ECD sequences were provided as paired AlphaFold2-Multimer v3 inputs on Colabfold^69^ 1.6.1, and 5 models were generated using 5 recycles for each. The top-ranked model for each pair was used to extract pLDDT, pTM, and ipTM scores for downstream comparison of structural confidence across pair groups.

### 3D network plot design and graph connectivity score estimation

To construct the 3D heatmap first a two dimensional Kawada-Kawaii spring network showing the enrichment connections of bait proteins for 49 high quality proximity labeling experiments was created. High quality refers to the enrichment of the targeted bait protein as one of the top hits in the experiment. The 2-dimensional bait network was then transformed to lay on a 3D sphere with radius = 1. This was done by finding a given node’s position and estimating theta and phi to create a new 3D spherical coordinate using the following equation:

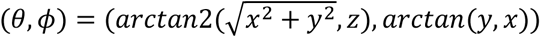

Pole points at 0, 0, 1 and 0, 0, -1 were added to the transformed points. For a given protein of interest, the normalized fold change for a given bait experiment was displayed on the corresponding bait node. A solid shell was then added to the 3D network and the scipy griddata function was then used to interpolate fold change values for the space between bait nodes. The graph connectivity score was then estimated by integrating over the fold change values on the sphere using the following equation:

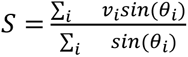

### Viral replication assay

Jurkat WT or IL10RB KO Jurkat cells were counted and resuspended at 200,000 cells/mL and seeded in 96 well plates at 20,000 cells per well in 100 uL of RPMI-1640 + 10% FBS + 1% pen/strep. 24 hour later, cells were stimulated with 0/1/10 ng/mL of IFNα2 (Thermo Scientific #300-02AA-20UG) in 5 uL of RPMI-1640. 24 hours later, cells were challenged with VSV-GFP pseudotyped with rVSV-GFP-RabiesV-G^70^ at an intended multiplicity of infection of 2 and incubated at 37 degrees for one hour before beginning imaging on an Incucyte S3 (Sartorius). Six replicate wells per genotype and IFN condition were imaged in both phase contrast and green fluorescence channels using default acquisition parameters on an Incucyte S3 (Sartorius). For image acquisition, 9 fields per well were captured at 20× magnification, and images were quantified using the corresponding analysis settings for each magnification.

### Protein structural analysis

All protein structures in this work were analyzed and generated in ChimeraX version 1.11.1^71^.

### General Information for Synthetic Procedures

Diazirine hydrochloride ((4-(3-(trifluoromethyl)-3H-diazirin-3-yl)phenyl)methanaminium chloride) was purchased from TCI and used without further purification. NHBoc-PEG3-CO2H (2,2-dimethyl-4-oxo-3,8,11,14-tetraoxa-5-azaheptadecan-17-oic acid) was purchased from Broadpharm and used without further purification. Iridium photocatalysts Ir-G3 ([Ir(dFCF3CO2Hppy)2dMebpy-DBCO]PF6)1 and Ir-G2 ([Ir(dFCF3ppy)2dMebpy-DBCO]PF6)2 were generous gifts from Ciaran Seath (UF Scripps). Biotin-PEG3-diazirine was prepared according to a modified previously reported route3 (see synthetic details and characterization data below). Solvents were purified with a PureSolv Solvent Purification System or used as commercially available anhydrous sure-seal bottles. Non-aqueous reagents were transferred under nitrogen via syringe. Organic solutions were concentrated under reduced pressure on a Büchi rotary evaporator using a water bath. Normal phase chromatographic purification of products was accomplished using forced-flow chromatography on a Biotage automated flash column chromatography system equipped with Biotage Sfär pre-packed silica gel columns (60 μm particle size). Thin-layer chromatography (TLC) was performed on Silicycle 0.25 mm silica gel F-254 plates. Visualization of the developed chromatogram was performed by UV lamp exposure and KMnO4 stain. 1H NMR spectra were recorded on a Bruker NEO-500 MHz and are internally referenced to residual protio CDCl3 (7.26 ppm) signals. CDCl3 was stored over K2CO3. Data for 1H NMR are reported as follows: chemical shift (δ ppm), integration, multiplicity (s = singlet, d = doublet, t = triplet, q = quartet, m = multiplet, dd = doublet of doublets, dt = doublet of triplets, br = broad), and coupling constant (Hz). Reversed-phase preparatory HPLC purification was carried out with an Agilent 1260 Infinity II system equipped with an Agilent preparatory HPLC column (5 μm, C18, 50 x 21.2 mm L x ID).

### Bioconjugation of iridium photocatalyst Ir-G2 to a secondary antibody

To a 0.5 mL microcentrifuge tube were added 20 uL of NHS-PEG4-azide (2.5 mM from a 5 mM stock solution in DMSO) and 20 uL of Ir-G2. The solution was incubated at room temperature in the dark for 4 hours. Into a 1.5 mL LoBind microcentrifuge tube was added 18 uL of the Ir-G2 solution (X uM, 45 eq.), 75 uL of goat anti-mouse antibody (6.67 uM from a 2 mg/mL or 13.3 uM stock, 1 eq.), and 57 uL of PBS. This sample was incubated at room temperature for 16 hours. A 40 kDa MWCO Zeba desalting spin filter (Thermo Fisher Scientific #87767) was then used to clean up the reaction solution. The photocatalyst-antibody conjugate was analyzed by BCA protein assay kit for total protein concentration and A350 for Ir concentration against serial dilutions of BSA and unconjugated Ir-G2 as standards, respectively to ensure a final concentration of 1.0 mg/mL and an Ir/primary Ab ratio of 6-8. The conjugate was then stored at 4°C and used within 7 days of conjugation.

### Bioconjugation of iridium photocatalyst Ir-G3 to a secondary antibody

To a 0.5 mL microcentrifuge tube were added 20 uL of NHS-PEG12-azide (2.5 mM from a 5 mM stock solution in DMSO) and 20 uL of Ir-G3. The solution was incubated at room temperature in the dark for 4 hours. Into a 1.5 mL LoBind microcentrifuge tube were added 18 uL of the Ir-G3 solution, 75 uL of goat anti-mouse antibody (6.67 uM from a 2 mg/mL or 13.3 uM stock), and 57 uL of PBS. This sample was incubated at room temperature for 16 hours. A 40 kDa MWCO Zeba desalting spin filter was then used to clean up the reaction solution. The photocatalyst-antibody conjugate was analyzed by BCA protein assay kit for total protein concentration and A350 for Ir concentration against serial dilutions of BSA and unconjugated Ir-G3 as standards, respectively to ensure a final concentration of 1.0 mg/mL and an Ir/primary Ab ratio of 6-8. The conjugate was then stored at 4°C and used within 7 days of conjugation.

### tert-butyl (3-oxo-1-(4-(3-(trifluoromethyl)-3H-diazirin-3-yl)phenyl)-6,9,12-trioxa-2-azatetradecan-14-yl)carbamate (NHBoc-PEG3-diazirine) synthesis

To a 20 mL scintillation vial, starting acid 2,2-dimethyl-4-oxo-3,8,11,14-tetraoxa-5-azaheptadecan-17-oic acid (NHBoc-PEG3-CO2H, 200 mg, 0.62 mmol), HATU (237 mg, 0.62 mmol, 1 equiv), and (4-(3-(trifluoromethyl)-3H-diazirin-3-yl)phenyl)methanaminium chloride (diazirine, 161 mg, 0.75 mmol, 1.2 equiv) were added before suspension in DMF (6 mL, 100 mM). To this mixture, diisopropylethylamine (DIPEA, 540 μL, 3.1 mmol, 5 equiv) was added. The reaction mixture was stirred in the dark at room temperature for 16 hours before combining with 30 mL H2O and extracting with EtOAc (4 x 10 mL). The combined organic fractions were then concentrated by rotary evaporation and purified by SiO2 gel column chromatography (0–20% MeOH/DCM) to give the title compound as a white solid (300 mg, 93% yield). 1H NMR (500 MHz, CDCl3) δ 7.32 (d, J = 8.2 Hz, 2H), 7.15 (d, J = 8.0 Hz, 2H), 6.96 (s, 1H), 4.92 (s, 1H), 4.45 (d, J = 6.0 Hz, 2H), 3.76 (t, J = 5.7 Hz, 2H), 3.66 – 3.57 (m, 4H), 3.55 – 3.48 (m, 4H), 3.46 (t, J = 5.2 Hz, 2H), 3.30 – 3.22 (m, 2H), 2.53 (t, J = 5.7 Hz, 2H), 1.43 (s, 9H).

### N-(3-oxo-1-(4-(3-(trifluoromethyl)-3H-diazirin-3-yl)phenyl)-6,9,12-trioxa-2-azatetradecan-14-yl)-5-((3aS,4S,6aR)-2-oxohexahydro-1H-thieno[3,4-d]imidazol-4-yl)pentanamide (biotin-PEG3-diazirine) synthesis

Boc-protected diazirine-PEG3-amine (300 mg, 0.58 mmol) was suspended in a 4.0 M HCl/dioxane solution (3 mL) in a 20 mL scintillation vial and stirred at room temperature in the dark for 1 hour. The reaction was then evaporated, the resulting residue redissolved in DMF (2.0 mL), and diisopropylethylamine added (300 μL, 1.7 mmol, 3 equiv). To a separate 20 mL scintillation vial biotin (280 mg, 1.2 mmol, 2 equiv) and HATU (440 mg, 1.2 mmol, 2 equiv) were added. To this vial DMF (2.0 mL) and diisopropylethylamine (200 μL, 1.2 mmol, 2 equiv) were added and the mixture stirred at room temperature for 15 minutes. The activated biotin solution was then added to the crude deprotected diazirine-PEG3-amine solution and stirred in the dark at room temperature for 16 hours. The crude reaction mixture was combined with 20 mL H2O and extracted with EtOAc (4 x 10 mL). The organic fractions were combined and washed with brine (3 x 10 mL), dried over MgSO4, and concentrated by rotary evaporation and purified by SiO2 gel column chromatography (0–50% MeOH/DCM). Product-containing fractions were then combined, concentrated, and subjected to preparative scale reversed-phase HPLC (C18, 0-80% MeCN/0.1%FA in H2O/0.1%FA) to give the title compound as a white solid (71 mg, 29% yield). 1H NMR (500 MHz, CDCl3) δ 7.33 (d, J = 8.2 Hz, 2H), 7.15 (d, J = 8.0 Hz, 3H), 6.39 (s, 1H), 5.87 (s, 1H), 4.88 (s, 1H), 4.52 – 4.43 (m, 3H), 4.30 – 4.26 (m, 1H), 3.78 (t, J = 5.8 Hz, 2H), 3.66 – 3.49 (m, 10H), 3.46 – 3.34 (m, 2H), 3.14 (td, J = 7.3, 4.5 Hz, 1H), 2.91 (dd, J = 12.8, 4.9 Hz, 1H), 2.71 (d, J = 12.8 Hz, 1H), 2.54 (t, J = 5.8 Hz, 2H), 2.19 (td, J = 7.2, 1.9 Hz, 2H), 1.67 (dtd, J = 24.4, 8.8, 6.4 Hz, 4H), 1.48 – 1.39 (m, 2H). 19F NMR (471 MHz, CDCl3) δ -65.25.

## Data availability

Raw data for proteomics files are publicly available on MassIVE with accession number MSV000096983 and on ProteomeXchange with accession number PXD060303. Processed proteomics results and data used to make all graphs included in main text and extended data figures are included as Source Data files.

## Code availability

Code used for analysis and visualization in this work is open source and freely available at https://github.com/brendanfloyd/cell_surface_map.

