## Extended Data figures for "A nanoscale map of the human cell surface proteome reveals spatial and functional organization"

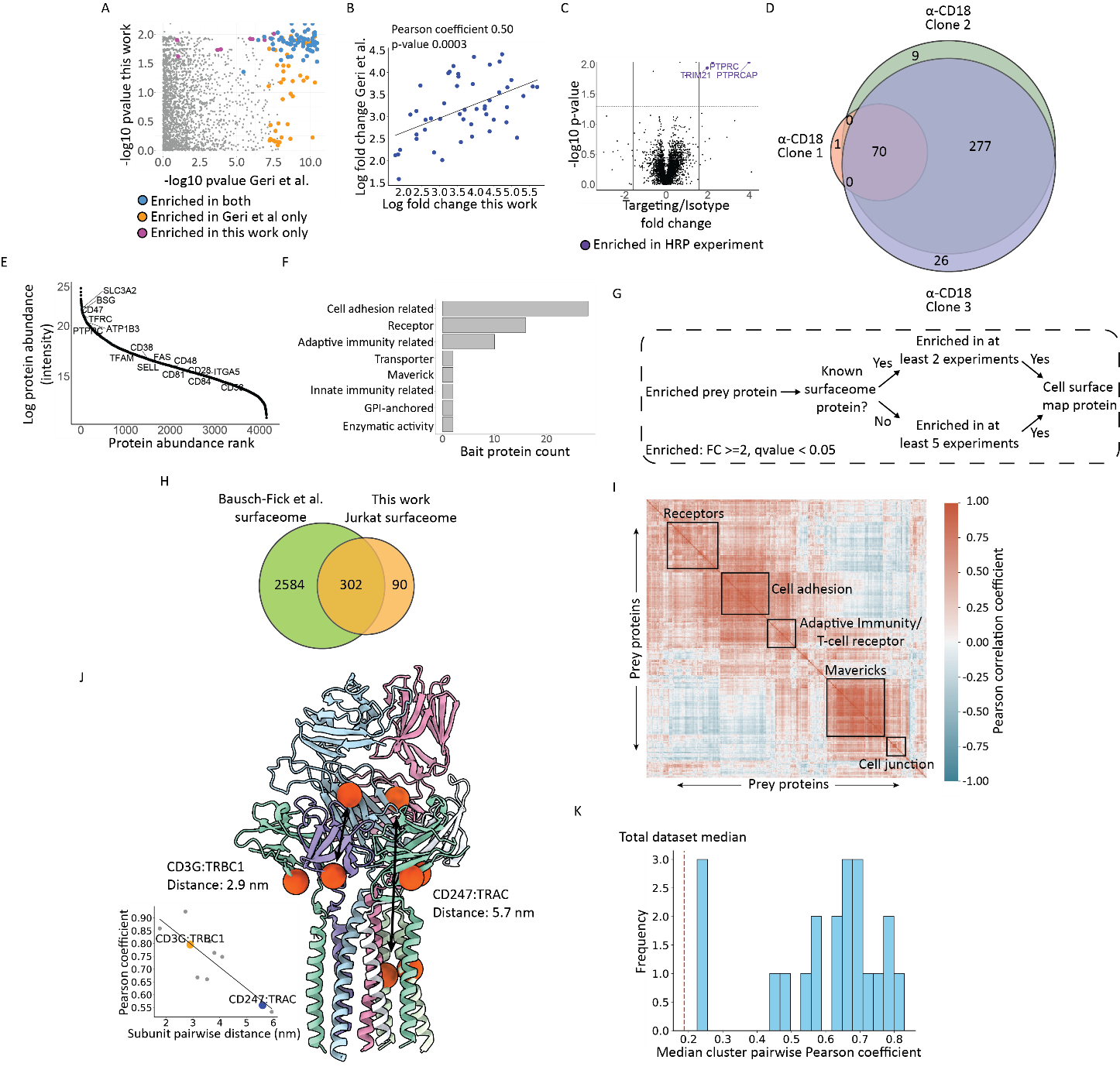

**Extended Data Figure 1** Validation of cell surface proximity labeling experiments collected in Jurkat cells and additional data for the prey-centric analysis of the Jurkat cell surface proteome. (A) Comparison of estimated FDR-corrected p-values from a CD45 targeting proximity labeling experiment to a previously published dataset. Proteins enriched (p-value < 0.05 and fold change ≥ 2). (B) Comparison of fold change values of enriched proteins from the CD45 proximity labeling experiment. The p-value was estimated using SciPy’s two-sided test of the null hypothesis that the true correlation coefficient is zero. (C) Volcano plot showing the results of a CD45-targeting µMap proximity labeling experiment. All enriched proteins were also enriched in the HRP proximity labeling experiment. P-values were estimated using a two-sided t-test with false discovery rate (FDR) with correction performed using permutation-based FDR with 250 randomizations for n = 3 biological replicates. (D) Venn diagram of enriched proteins (p-value < 0.05 and fold change ≥ 2) from an HRP proximity labeling experiment targeting CD18/ITGB2 with 3 different antibody clones. (E) Rank-abundance curve from a WGA-HRP surfaceomics experiment with proteins chosen as bait proteins for proximity labeling experiments highlighted. (F) Bar plot showing the different functional groups for the 65 bait proteins used for proximity labeling on Jurkat cells. (G) Workflow for determining if a protein is a cell surface protein expressed on Jurkat cells. (H) Venn diagram comparison of the estimated Jurkat surfaceome for this work compared to a published machine learning-predicted surfaceome. (I) Hierarchically clustered heatmap of Pearson correlation values estimated from the fold change enrichment of all prey proteins on the Jurkat cell surface proteome. Enriched functional annotations are highlighted for the larger clusters of functional units. (J) Structure of the TCR-CD3 protein complex with the center of mass of each subunit highlighted as an orange sphere. Pairwise distance measurements were taken from the center of mass between two subunits and compared to the estimated pairwise Pearson correlation coefficient values. Two examples of pairwise measurements are highlighted on the structure and the scatterplot. (K) Distribution of median pairwise Pearson coefficients for the clusters shown in Figure 1F. The median value for pairwise Pearson coefficients for the entire Jurkat surfaceome is shown as a red line.

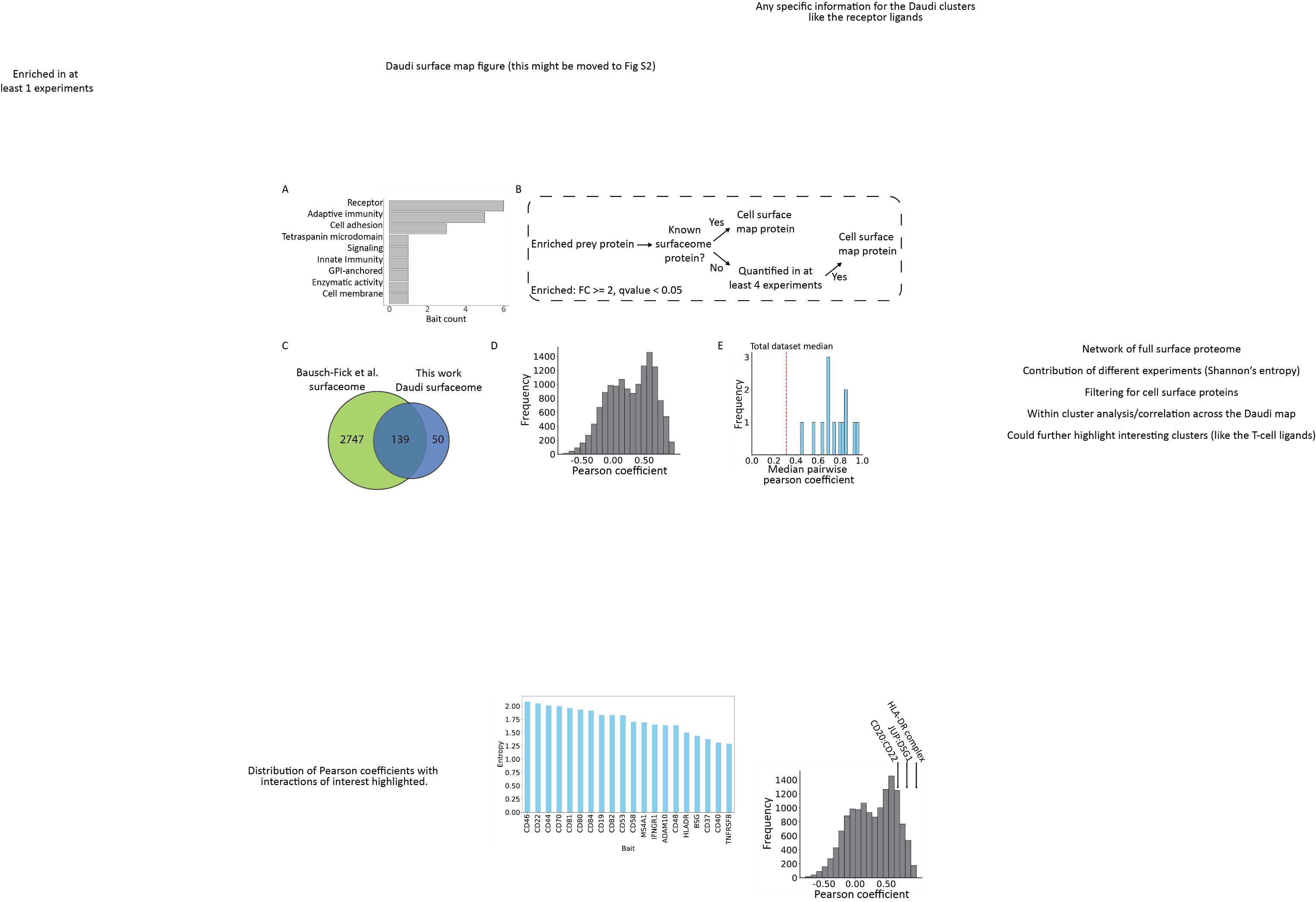

**Extended Data Figure 2** Characterization of cell surface protein characteristics on Daudi cell surfaces. (A) Bar plot showing the different functional annotations for the 20 bait proteins used for proximity labeling on Daudi cells. (B) Workflow for determining if a protein is a cell surface protein expressed on Daudi cells. (C) Venn diagram comparison of the estimated Daudi surfaceome for this work compared to a published machine learning-predicted surfaceome. (D) Distribution of all pairwise Pearson correlation coefficients for Daudi cell surface proteins. (E) Distribution of median pairwise Pearson coefficients for the clusters shown in Figure 2G. The median value for Pearson pairwise coefficients for the entire Daudi surfaceome is shown as a red line.

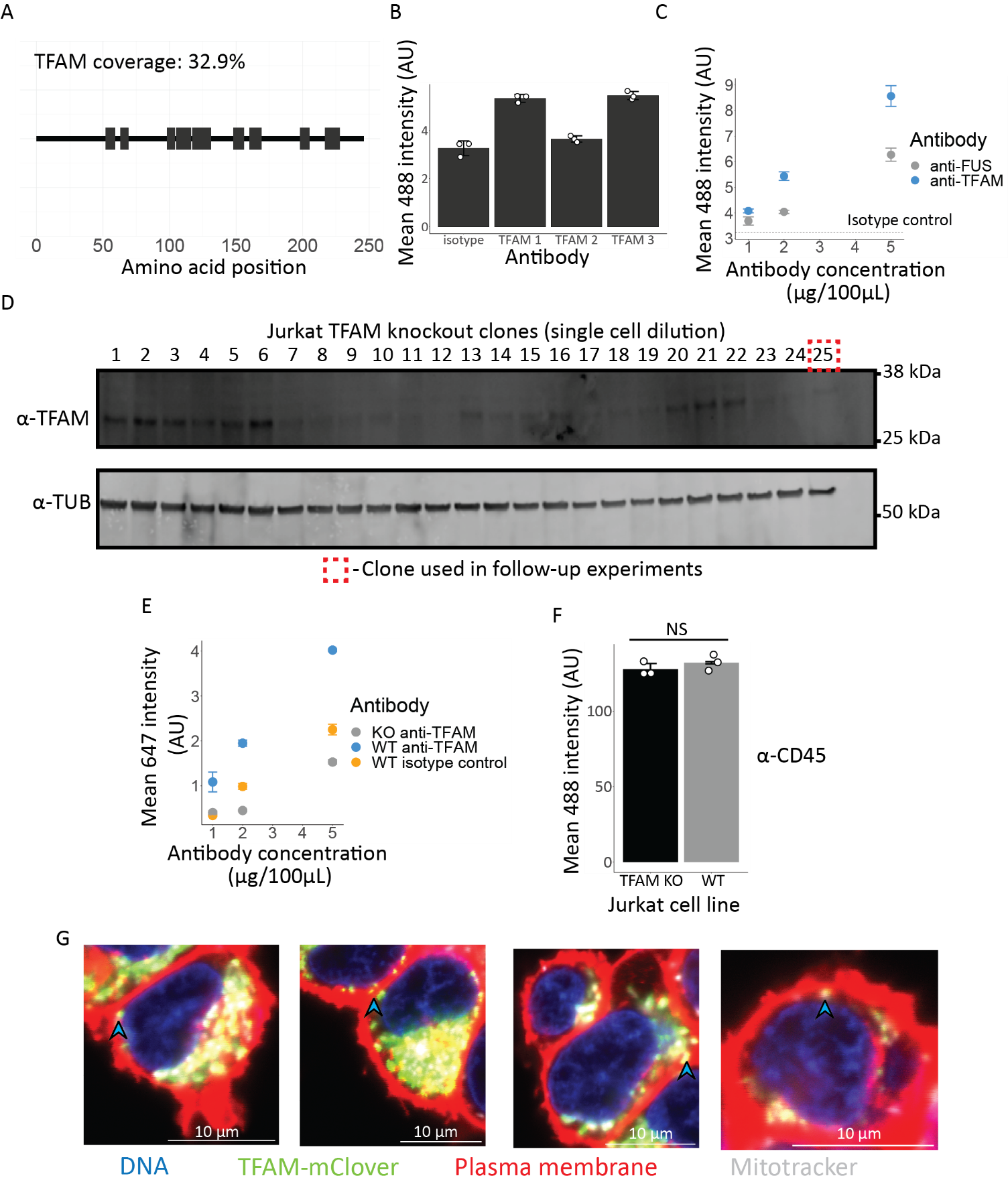

**Extended Data Figure 3** Validation and characterization of a maverick cell surface protein. (A) Peptide coverage of TFAM from proximity labeling experiments. The grey bars correspond to peptides detected by mass spectrometry. (B) Bar plot showing the flow cytometry results for proteins stained on the surface with 3 different α-TFAM antibodies and an isotype control. Error bars correspond to standard deviation from the mean. α-TFAM 3 (Proteintech) was used in all subsequent experiments. (C) Flow cytometry results from cell surface staining with an α-TFAM and a previously validated α-FUS antibody at 3 different concentrations. The isotype control binding is shown as a dashed line and error bars correspond to standard deviation from the mean. (D) Western blot results following CRISPR-Cas9 knockout of TFAM in Jurkat cells and single-cell dilution to isolate clones with TFAM knocked out. Clone 25 was used in all subsequent experiments. (E) Flow cytometry results from cell surface staining with an anti-TFAM or isotype control antibody at 3 different concentrations on wildtype (WT) and TFAM knockout (KO) Jurkat cells. Error bars correspond to the standard deviation of the mean. (F) Flow cytometry results from cell surface staining with an anti-CD45 antibody on Jurkat wildtype and TFAM knockout cells. Student’s t-test p-value of 0.332 and t-statistic of -1.103 on n = 3 biological replicates. Error bars correspond to the standard deviation from the mean. (G) Additional confocal microscopy image of HEK-293T TFAM-mClover (green) expressing cell lines stained with a CellMask Orange plasma membrane stain (red), Hoescht for nuclear DNA (blue), and Mitotracker red (white). mClover fluorescence was detected in the 488 nm channel, the plasma membrane stain in the 561 nm channel, Hoechst in the 405 nm channel, and MitoTracker Red in the 647 nm channel. Images are representative of an experiment independently repeated 3 times.

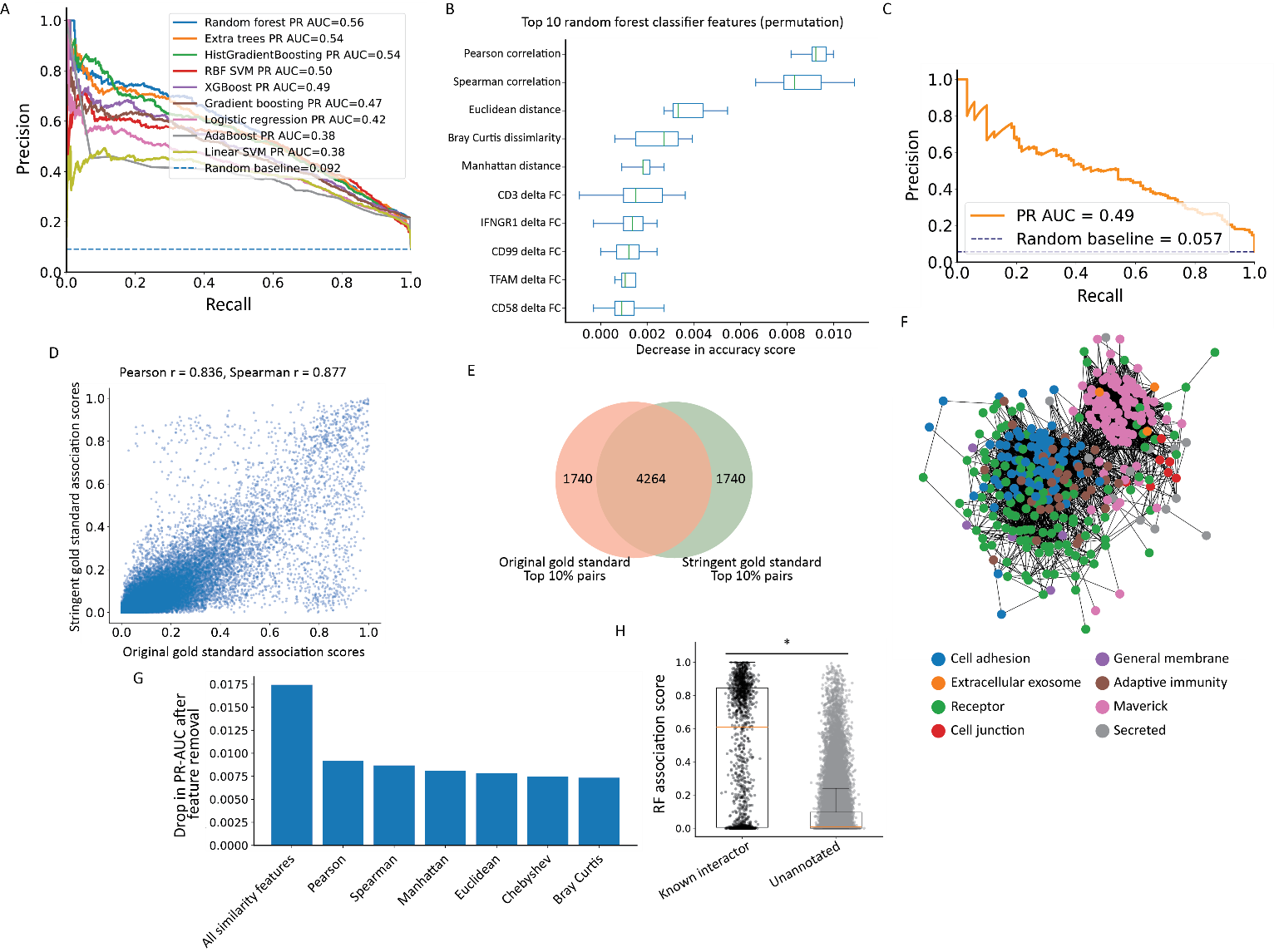

**Extended Data Figure 4.** Additional information on the random forest classifier for prioritizing cell surface associations. (A) Precision-recall curves comparing the random forest classifier to eight additional supervised machine learning classifiers for prioritizing gold-standard cell surface protein associations from the proximity labeling-derived feature matrix. Classifier performance was evaluated using stratified 5-fold cross-validation with pooled out-of-fold prediction scores. Precision-recall area under the curve (PR-AUC) values are shown in the legend for each classifier, and the dashed line indicates the random baseline determined by the fraction of positive labels in the benchmark set. (B) The top 10 most important random forest classifier features of the 70 used for predicting cell surface associations on Jurkat cells from a permutation test. For the boxplot the central line corresponds to the median and the whiskers extend to the most extreme data points within 10 times the interquartile range (IQR). (C) Precision-recall curve for a random forest classifier trained using a more stringent gold-standard positive set incorporating UniProt PPIs and BioGRID multi-validation (BioGRID-MV) physical interaction pairs (D) Comparison of random forest association scores generated using the original curated gold-standard PPI set and the more stringent gold-standard PPI set incorporating UniProt PPIs and BioGRID-MV physical interaction pairs. Each point represents a protein pair scored by both classifiers. (E) Venn diagram comparing the top 10% highest-scoring protein pairs from random forest classifiers trained with the original curated gold-standard PPI set or the more stringent gold-standard PPI set incorporating BioGRID-MV physical interaction pairs. (F) Kawada-Kawaii network where each node is a Jurkat cell surface protein, and the edges are drawn between nodes with an association score that passed a 10% false-discovery rate threshold. Nodes are colored according to the functional association of their assigned cluster shown in Figure 1F. (G) Effect of similarity feature ablation on classifier performance, shown as the decrease in PR-AUC after removing individual similarity features or all similarity features prior to random forest training. (H) Random forest association scores for known interacting protein pairs and unannotated protein pairs. Mann-Whitney U test p-value = 3.61 × 10^-281^. For the known interactor category n = 1,547 and for the unannotated category n = 58,484.

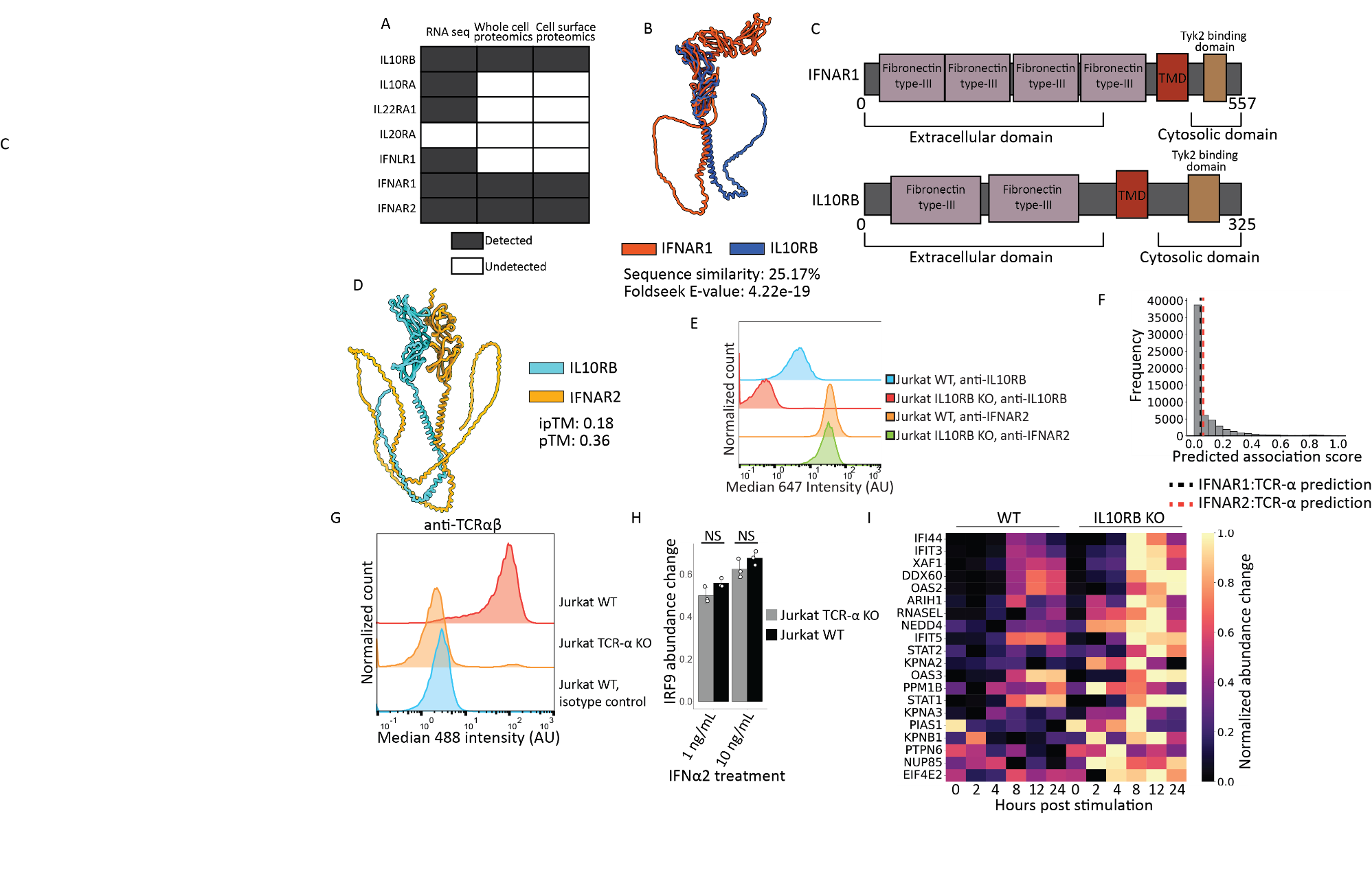

**Extended Data Figure 5** Characterization of IL10RB as a type I interferon regulator. (A) Identification of IL10RB interactors in Jurkat cells by RNA-seq, whole cell proteomics, and cell surface proteomics. RNA-seq data is from Felce et al. (2021). (B) Aligned AlphaFold 3 predicted structures of IFNAR1 and IL10RB which share structural similarity as read out using Foldseek. (C) Protein domain architecture of IFNAR1 and IL10RB. (D) Top AlphaFold 3 prediction between IFNAR2 and IL10RB with ipTM and pTM values of 0.18 and 0.36. (E) Flow cytometry intensity distributions for Jurkat wild type (WT) and IL10RB knockout (KO) cells stained with an anti-IL10RB or anti-IFNAR2 antibody. A secondary antibody conjugated with Alexa Fluor 647 was used for fluorescent readout. (F) Distribution of predicted association scores for all 60,031 predictions with the prediction for TCR-α and the two subunits of the type I interferon protein complex highlighted. The association scores for TCR-α:IFNAR1 and TCR-α:IFNAR2 are 0.05 and 0.06 respectively. (G) Flow cytometry intensity distributions for Jurkat wild type (WT) and TCR-α knockout (KO) stained with an anti-TCR-αβ or isotype control antibody. (H) Intracellular flow cytometry measurement of cytosolic IRF9 abundance change in IFNα2 stimulated Jurkat wild type (black) and TCR-α knockout lines (grey) at IFNα2 concentrations of 1 and 10 ng/mL. Two-sided Student’s t-test p-values at the different concentrations are 0.078 and 0.176 (t-statistics of 2.36 and 1.64) for 1 and 10 ng/mL respectively with n = 3 biological replicates for each. Error bars correspond to the standard deviation of the mean. (I) Heatmap showing the 20 interferon stimulated genes (ISGs) with the largest stimulation changes in the Jurkat IL10RB knockout (KO) line in comparison to the wild type line when stimulated with IFNα2 at various time points.

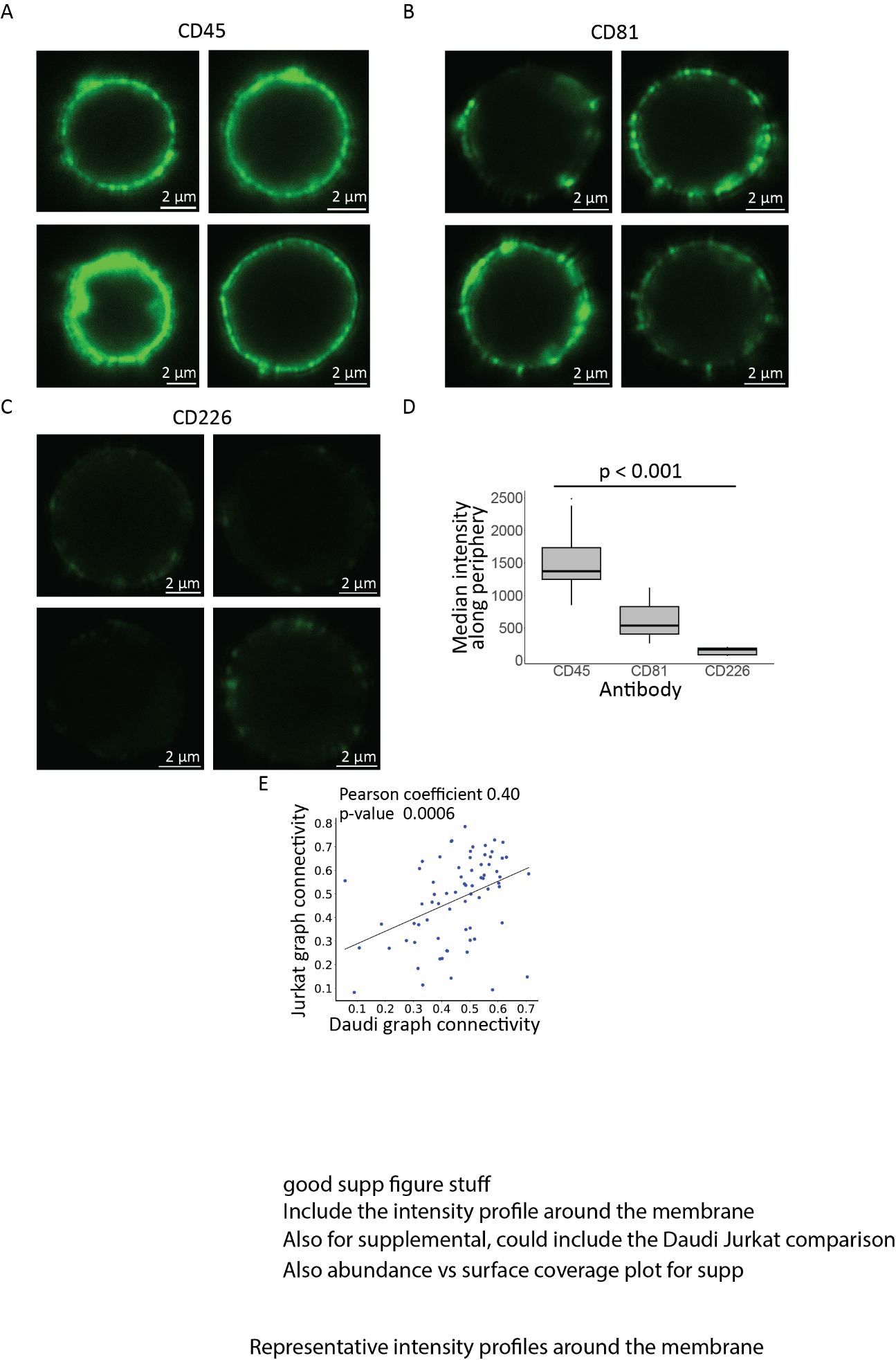

**Extended Data Figure 6** Additional data on the graph connectivity score and cell surface staining for confocal microscopy. (A) Additional representative images for cells stained with an anti-CD45 primary antibody and an Alexa Fluor 488 secondary antibody. (B) Additional representative images for cells stained with an anti-CD81 primary antibody and an Alexa Fluor 488 secondary antibody. This experiment was repeated independently 3 times. (C) Additional representative images for cells stained with an anti-CD226 primary antibody and an Alexa Fluor 488 secondary antibody. (D) Box plot showing the median intensity along the cell periphery for cells stained with anti-CD45, anti-CD81, and anti-CD226 antibodies. A one-way ANOVA was done to determine differences in the mean with an F-statistic of 32.328 and p-value of 1.549e-7 for n = 10 cells for CD45 and CD81 samples and n=7 cells for CD226 samples. The whiskers extend to the smallest and largest data points within 1.5 times the interquartile range from the first and third quartiles, respectively. The box is centered on the median value of the distribution. (E) Scatterplot showing the estimated graph connectivity for antigens both on Jurkat and Daudi cells. The line represents the line of best fit for the data and the p-value was estimated using SciPy’s two-sided test of the null hypothesis that the true correlation coefficient is zero.
